# Structural and functional analysis of the minimal orthomyxovirus-like polymerase of Tilapia Lake Virus from the highly diverged *Amnoonviridae* family

**DOI:** 10.1101/2023.08.14.553164

**Authors:** Benoit Arragain, Martin Pelosse, Albert Thompson, Stephen Cusack

## Abstract

Tilapia Lake Virus (TiLV), a recently discovered pathogen of tilapia fish, belongs to the *Amnoonviridae* family from the *Articulavirales* order. Its ten genome segments have characteristic conserved ends and encode proteins with no known homologues, apart from the segment 1, which encodes an orthomyxo-like RNA-dependent-RNA polymerase core subunit. Here we show that segments 1-3 encode respectively the PB1, PB2 and PA-like subunits of an active heterotrimeric polymerase that maintains all domains found in the distantly related influenza polymerase, despite an unprecedented overall size reduction of 40%. Multiple high-resolution cryo-EM structures of TiLV polymerase in pre-initiation, initiation and active elongation states, show how it binds the vRNA and cRNA promoters and performs RNA synthesis, with both transcriptase and replicase configurations being characterised. However, the highly truncated endonuclease-like domain appears inactive and the putative cap-binding domain is autoinhibited, emphasising that many functional aspects of TiLV polymerase remain to be elucidated.

## Introduction

Tilapia Lake Virus (TiLV) was first isolated from moribund fish following mass die offs of Nile Tilapia (*Oreochromis niloticus*) in fish farms in Israel^1^. Subsequent analysis showed that it is an enveloped, negative-sense single-stranded RNA virus with ten genome segments, each with at least one open-reading frame (ORF)^2^. Segment 1 encodes a protein of 519 residues containing motifs characteristic of the orthomyxovirus RNA-dependent RNA polymerase (RdRp) PB1 subunit, whereas the other nine ORFs exhibit no sequence homology to any other known protein. This, together with the presence of conserved, quasi-complimentary 5′ and 3′ termini on each segment led to the tentative classification of TiLV as a novel orthomyxo-like virus^2^. TiLV outbreaks have been reported in at least 16 countries located on four continents. The severity of TiLV outbreaks varies from asymptomatic with no associated mortality to over 90% mortality (Pulido *et al*, 2019, https://doi.org/10.1016/j.aquaculture.2019.04.058)^3, 4^. TiLV therefore poses a significant threat to the global tilapia aquaculture industry, which provides proteinaceous food to millions of people, especially in the developing world.

TiLV is now defined as a new species, *Tilapia tilapinevirus*, in the Tilapinevirus genus, within the *Amnoonviridae* family, which together with the *Orthomyxoviridae*, form the *Articulavirales* order (https://ictv.global/taxonomy/taxondetails?taxnode_id=202206024). Whilst the phylogeny of different isolates^5^, detection and pathogenesis of the virus and possible treatments by vaccination^6^ or drugs^7^ are now being extensively studied, the molecular virology of TiLV is poorly understood, partly due to the current lack of reverse genetics or minigenome systems^8^. In addition, the uniqueness of the putative TiLV encoded proteins, apart from a recognisable polymerase PB1-like subunit, precludes hypotheses as to protein function based on sequence homology or structure prediction using Alphafold^9^.

Here we set out to structurally characterize the TiLV encoded replication machinery, which should comprise at least an RdRp, potentially heterotrimeric and a nucleoprotein. We used co-expression of all TiLV ORFs in insect cells to identify proteins co-purifying with the segment 1 protein, the putative core RdRp subunit. This enabled purification of a heterotrimeric complex comprising proteins from segments 1-3, which could be structurally characterised by high-resolution cryogenic electron microscopy (cryo-EM) (Table 1). We show that this heterotrimer resembles the influenza virus polymerase (FluPol), with segments 2 and 3 being respectively PB2- and PA-like in domain structure and fold, even though the TiLV polymerase (155 kDa) is remarkably only 57% of the overall molecular FluPol mass (270 kDa). Recombinant TiLV polymerase specifically binds the conserved 5′ and 3′ genome and anti-genome ends, with mode A and mode B promoter structures similar to other segmented negative strand RNA viruses^10–14^. Furthermore, it is active in RNA synthesis with an appropriate primer. Our structural and functional characterisation of TiLV polymerase is a first step towards a detailed understanding of the transcription/replication machinery of this intriguing virus.

**Table 1.** Summary of TiLV polymerase structures.

| Structure No. | Short name | Resolution | RNA/promoter mode | Description | PDB/EMDB |
| --- | --- | --- | --- | --- | --- |
| **vRNA initiation** |  |  |  |  |  |
| 1 | Full vRNA initiation with CTP | 2.73Å | vRNA/A | Full mode A structure with 3' end in the active site and incoming CTP at +1 | PDB 8PSN<br>EMD-17857 |
| 2 | Core vRNA initiation with CTP | 2.40Å | vRNA/A | Core only mode A structure with 3' end in the active site and incoming CTP at +1 | PDB 8PSO<br>EMD-17858 |
| **cRNA pre-initiation** |  |  |  |  |  |
| 3 | Core cRNA pre-initiation mode A | 2.65Å | cRNA/A | Core only mode A structure with 3' end in the active site | PDB 8PSQ,<br>EMD-17860 |
| 4 | Core with endo cRNA pre-initiation mode A | 2.80Å | cRNA/A | Core with endo, mode A structure with 3' end in the active site | PDB 8PT7<br>EMD-17869 |
|  | **cRNA mode B** |  |  |  |  |
| 5 | Closed core with endo cRNA mode B | 2.83Å | cRNA/B | Core with endo, mode B structure with 3' end in the secondary site (weak duplex) | PDB 8PSS<br>EMD-17861 |
| **vRNA pre-initiation** |  |  |  |  |  |
| 6 | Core vRNA pre-initiation mode A | 3.18Å | vRNA/A | Core, mode A structure with 3' end in the active site | PDB 8PSU<br>EMD-17862 |
| **vRNA elongation** |  |  |  |  |  |
| 7 | Full vRNA elongation with CpNHpp | 2.96Å | vRNA | Full elongation structure stalled with CpNHpp at +1 | PDB 8PSX<br>EMD-17864 |
| 8 | Full vRNA elongation with CpNHpp and additional mode B promoter | 2.42Å | vRNA/B | Full elongation structure stalled with CpNHpp at +1 with additional mode B promoter | PDB 8PSZ<br>EMD-17865 |
| **vRNA mode B** |  |  |  |  |  |
| 9 | Full with open core vRNA mode B | 2.59Å | vRNA/B | Full mode B structure with 3' end in the secondary site (strong duplex) | PDB 8PT2<br>EMD-17866 |
| 10 | Open core with rotated endo-NLS vRNA mode B | 2.73Å | vRNA/B | Core with rotated Endo-NLS mode B structure (open, strong duplex) | PDB 8PTH<br>EMD-17871 |
| 11 | Closed core with rotated endo-NLS vRNA mode B | 2.86Å | vRNA/B | Core with rotated Endo-NLS mode B structure (closed, weak duplex) | PDB 8PTJ<br>EMD-17872 |
| **Replicase** |  |  |  |  |  |
| 12 | Replicase initiation with CpNHpp | 2.90Å | vRNA/B | Core with rotated Endo-NLS and CBD (replicase conformation) mode A with 3' end in the active site and incoming CpNHpp at +1 | PDB 8PT6<br>EMD-17868 |

## Results

### Identification of the Tilapia Lake Virus polymerase

TiLV (*Amnoonviridae* family) segment 1 ORF contains conserved orthomyxo-like RdRp motifs suggesting that it is a PB1-like subunit^2^. To identify other viral proteins that can interact with the segment 1 protein we added a purification tag at its C-terminus and co-expressed it in insect cells together with the nine other putative TiLV proteins using the EMBacY bacmid (see Methods and Extended Data Fig. 1). After an initial affinity purification step, segment 1 protein (57 kDa) co-eluted with two other proteins, matching the molecular weights of segments 2 (51 kDa) and 3 (47 kDa), suggesting formation of a heterotrimeric complex. However, this initial complex was unstable upon further purification, likely due to suboptimal positioning of the purification tag (see Methods and Extended Data Fig. 1). A new expression construct encoding segment 1, 2 and 3 proteins, with segment 2 harbouring a TEV-cleavable deca-histidine purification tag on its C-terminus, enabled pure and stable heterotrimeric complex, suitable for further structural and functional studies, to be obtained (Fig. 1a).

**Figure 1.**
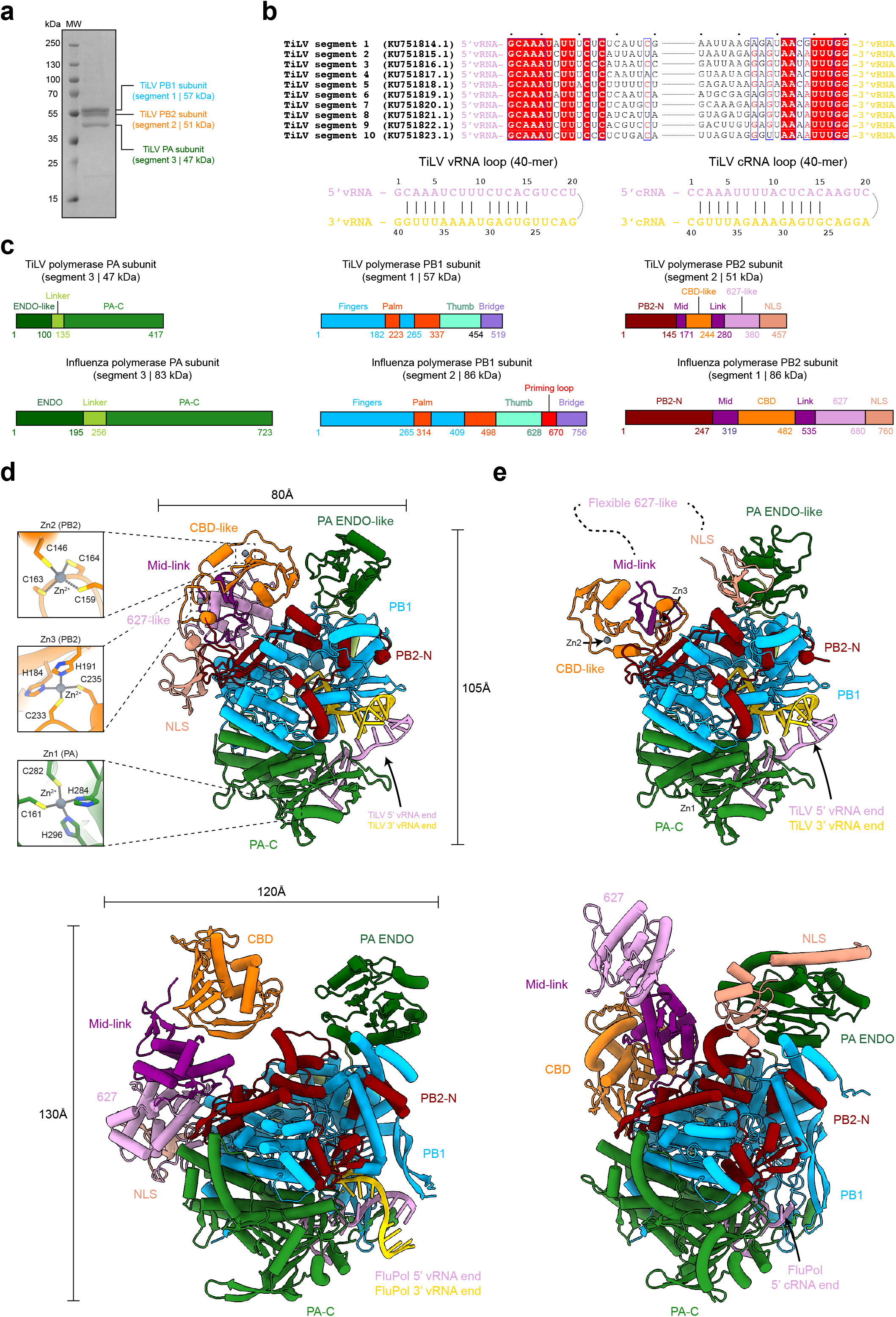
TiLV polymerase purification, promoter characterization and structural comparison with influenza polymerase. **a.** SDS-PAGE analysis of purified TiLV polymerase heterotrimeric complex. The molecular ladder (MW) is shown on the left side of the gel. TiLV polymerase subunits are indicated on the right side of the gel with PB1 (segment 1, 57 kDa), PB2 (segment 2, 51 kDa) and PA (segment 3, 47 kDa) subunits respectively coloured in blue, orange and green. **b.** Sequence alignment of the 20 first 5’ and 3’ end nucleotides from each TiLV genome segment. Corresponding GenBank numbers are indicated. Derived vRNA and cRNA loops (40-mer) from the segment 9 are shown with 5’ ends in pink and 3’ ends in gold. **c.** Schematic comparison of the domain structures of TiLV polymerase and FluPol subunits. Rectangle sizes are proportional for each corresponding polymerase domain. **d.** Cartoon representation of TiLV polymerase and FluPol (PDB: 4WSB) in the transcriptase conformation with respective domains coloured as in **c**. The three TiLV zinc-binding sites (Zn1, Zn2, Zn3) and their coordinating residues are displayed on the left. 5’ and 3’ vRNA ends are coloured as in **b**. TiLV polymerase and FluPol are aligned with each other using PB1 core as reference. **e.** Cartoon representation of TiLV polymerase and FluPol (PDB: 5EPI) in the replicase conformation, coloured as in **c** and **d**. TiLV PB2 627-like domain is flexible. TiLV polymerase and FluPol are aligned using PB1 core as reference.

### Reconstitution of TiLV polymerase with the promoter and cryo-EM structure determination

Viral polymerases from segmented negative stranded RNA viruses (sNSVs) typically interact directly with the conserved and quasi-complementary 5’ and 3’ viral genome ends, usually referred to as the promoter^11, 13, 14^. We therefore sought to identify TiLV promoter regions to be able to reconstitute an active TiLV heterotrimeric complex *in vitro*. Unfortunately, the previously highlighted viral promoter regions for each segment^2^ do not correspond to those deduced from the deposited negative sense RNA sequences. The correct 5’ and 3’ vRNA and complementary (cRNA) sequences for each TiLV segment show some features distinct from classical orthomyxoviral promoter sequences (Fig. 1b). The TiLV 5’ and 3’ ends are quasi-complementary over 15 nucleotides and, like bunyavirus promoters, are of the same length, without the bulge seen in the panhandle representation of the influenza promoter. Exceptionally, there is no complementarity between the first nucleotides of the 5’ and 3’ ends. We next synthesized 40-mer vRNA and cRNA loops, each corresponding to the joining of the first and last 20 nucleotides of the 5’ and 3’ vRNA/cRNA ends of segment 9 (Fig. 1b). Using single particle cryo-EM, we solved multiple sub-3Å resolution structures of reconstituted TiLV polymerase-promoter complexes that allowed unambiguous model building of the complete polymerase and promoter structures, in different functional states (Table 1, Extended Data Table 1, Supplementary Information 1-4).

### Overall structure of TiLV polymerase and comparison with influenza polymerase

TiLV polymerase is composed of PB1-, PB2- and PA-like subunits, encoded by segments 1-3 respectively. The heterotrimer is only 57% of the molecular weight of the distantly related orthomyxovirus FluPol (155 kDa versus 270 kDa). Remarkably, each subunit maintains the same domains as found in FluPol subunits, but each is minimised in size, except the RdRp core (Fig. 1c). The unusual presence of three zinc ions denoted Zn1, Zn2 and Zn3 may be one way by which domains are stabilized within the overall smaller architecture of TiLV polymerase. Zn1 is located in the PA C-terminal domain (PA-C), while Zn2 and Zn3 are located in the cap-binding domain-like (CBD-like) of PB2 (Fig. 1d). Furthermore, TiLV polymerase can take up the two functionally distinct overall configurations that are analogous to the FluPol transcriptase^15, 16^ and replicase^17–20^ conformations (Fig. 1de). Due to the structural relatedness of the TiLV and influenza polymerases, despite high sequence divergence, we henceforth drop the ‘-like’ qualification in future reference to TiLV polymerase subunits and domains, except for endonuclease-like (ENDO-like) domain and CBD-like domain, whose functionality remains to be clarified (see below).

### TiLV PA subunit

Comparison of the TiLV and FluPol PA subunits reveals that despite sharing no sequence homology and being 45% smaller (417 residues versus 723 residues), the structural fold and organization are conserved (Fig. 2a). Like FluPol, TiLV PA subunit is divided into three domains: a minimal N-terminal ENDO-like domain (1-100, 100 residues, 196 in FluPol); a short linker domain (101-135, 35 residues, 61 in FluPol); and the PA-C domain (136-417, 282 residues, 467 in FluPol) (Fig. 2a). TiLV ENDO-like domain is a highly truncated version of FluPol endonuclease and does not display any residues that could be involved in nuclease activity (Fig. 2b). Indeed, FluPol residues H41, E80, D108, E119 that coordinate divalent metal ions and K134, which performs nucleophilic attack, correspond to F5, Y29, D43, E56 and R62 respectively in the TiLV ENDO-like domain (Fig. 2c). *In vitro* endonuclease activity assays revealed that TiLV polymerase, in the presence of various divalent metal ions is unable to cleave single stranded RNA, whether or not bound to the 5’ and 3’ vRNA ends, in contrast to FluPol (Fig. 2d). The inactivity of TiLV ENDO-like domain is reminiscent of Thogoto and Dhori virus polymerases, whose ENDO-like domains also have non-functional active sites^21^, raising the questions as to if and how TiLV performs cap-dependent transcription and the functional role of the degenerate domain (Extended Data Fig. 2).

**Figure 2.**
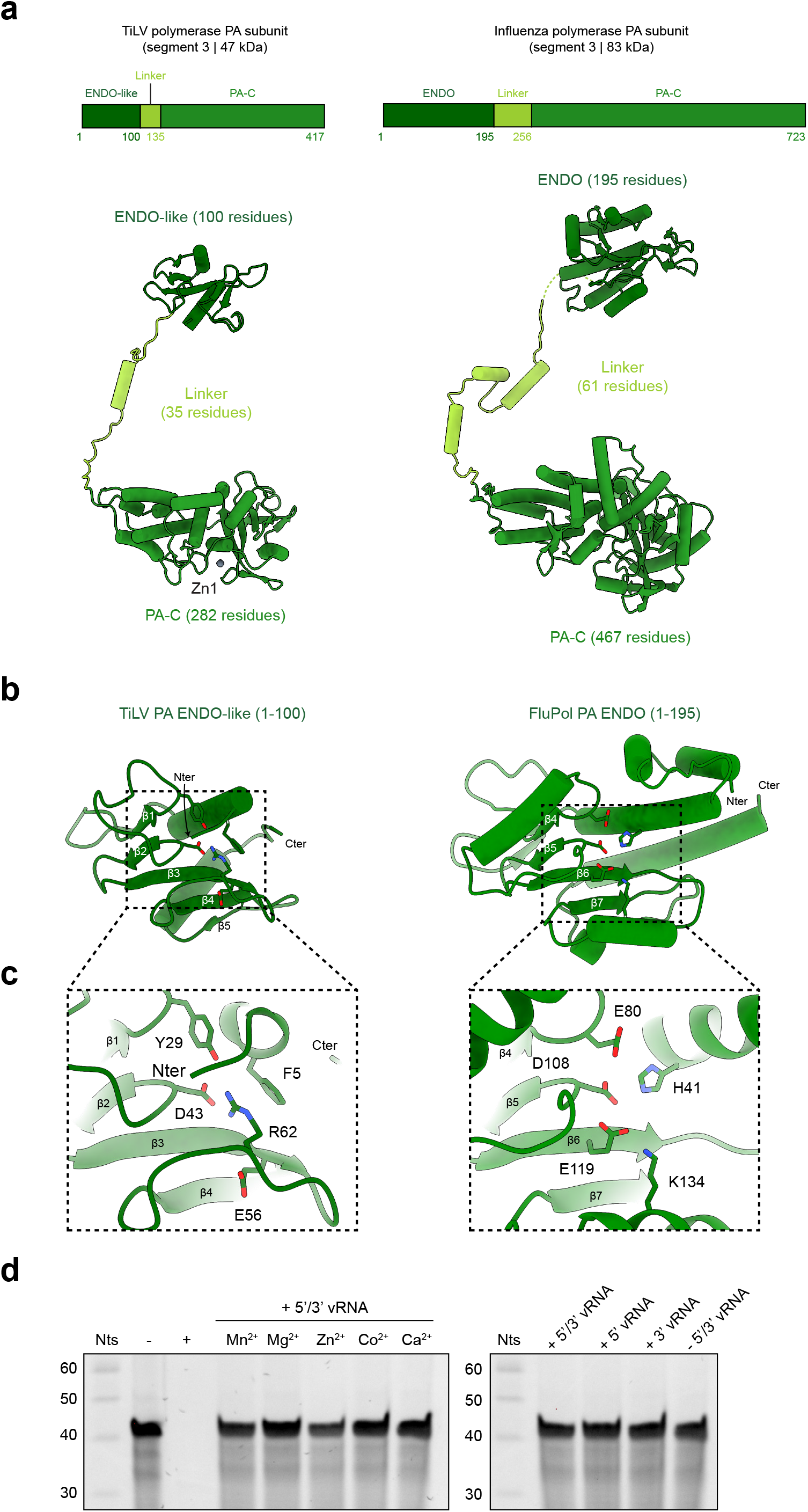
TiLV polymerase PA-like subunit in transcriptase conformation. **a.** Schematic and cartoon representations of TiLV polymerase and FluPol PA subunits extracted from their respective transcriptase conformation. Domains are coloured as in Fig. 1c. **b.** Comparison of TiLV PA ENDO-like (1-100) and FluPol PA ENDO (1-195) domain structures. The N-(N_ter_) and C-terminus (C_ter_) are indicated. The conserved β-sheets of TiLV polymerase and FluPol were used for domain alignment. **c.** Close-up of TiLV ENDO-like and FluPol ENDO domain active site. FluPol ENDO catalytic residues are displayed as well as the corresponding residues of TiLV ENDO-like domain. **d.** *In vitro* TiLV PA ENDO-like domain RNA cleavage activity. **Left:** Urea-PAGE analysis of the effect of different divalent metal ions on TiLV polymerase PA ENDO-like cleavage activity. Lanes “-” and “+” correspond to negative (no protein) and positive (A/H7N9 FluPol) controls. TiLV polymerase bound to the 5’ (1-15) and 3’ vRNA (1-15) ends, incubated with manganese (Mn^2+^), magnesium (Mg^2+^), zinc (Zn^2+^), cobalt (Co^2+^), or calcium (Ca^2+^), is not able to cleave single-stranded RNA. The decade marker (Nts) is shown on the left of the gel. The uncropped gel is provided as source data file. **Right:** Urea-PAGE analysis of the effect of vRNA ends on TiLV ENDO-like cleavage activity. TiLV polymerase bound to (i) the 5’ (1-15) and 3’ vRNA (1-15) ends (+5’/3’ vRNA), (ii) the 5’ vRNA end only (+5’ vRNA), (iii) the 3’ vRNA end only (+3’ vRNA) or (iv) no RNA (-5’/3’ vRNA) using manganese (Mn^2+^), is not able to cleave single-stranded RNA. The decade marker (Nts) is shown on the left of the gel. The uncropped gel is provided as source data file.

### TiLV PB1 subunit

TiLV PB1 subunit (57 kDa, 515 residues) encompasses the expected RdRp sub-domains, the fingers (1-182, 224-265), palm (183-223, 266-337), thumb (338-454) and bridge (455-515). They are structurally analogous to the FluPol PB1 domains, but are overall 44% smaller (Fig. 3a). TiLV PB1 structural minimization examples are numerous. The FluPol PB1 α-helices (84-123, 40 residues) are replaced by extended chain in TiLV PB1 (46-73, 28 residues) (Fig. 3b), the FluPol PB1 β-ribbon (162-222, 61 residues) is completely absent in TiLV PB1 (107-137, 31 residues) (Fig. 3c). More strikingly, the FluPol PB1 priming loop, which is an essential structural element in the orthomyxovirus genome replication initiation mechanism^22^, is completely absent in TiLV PB1 (Fig. 3d).

**Figure 3.**
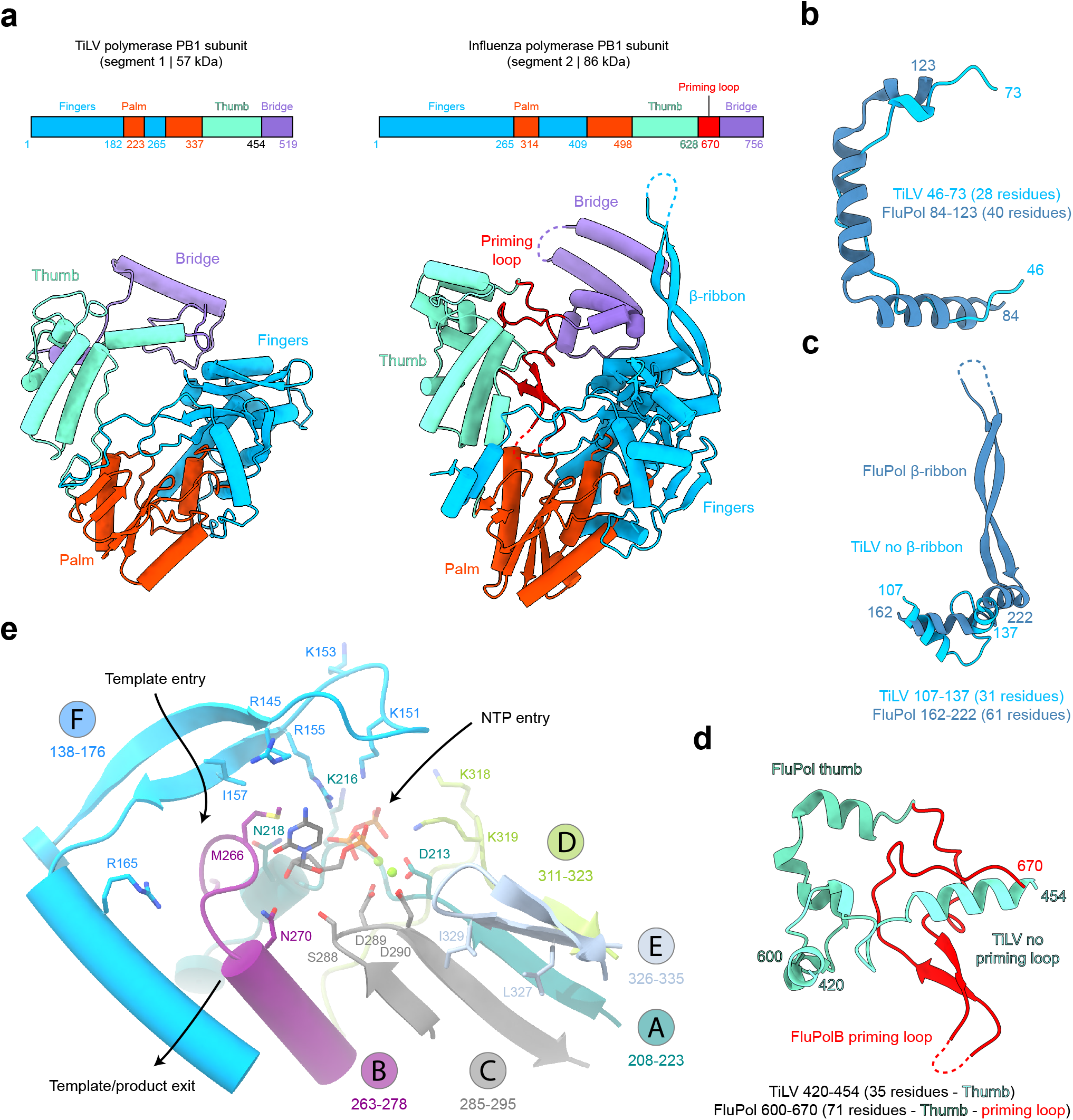
TiLV polymerase PB1 subunit. **a.** Schematic and cartoon representation of TiLV polymerase and FluPol PB1 subunits. Domains are coloured as in Fig. 1c. **b.** Miniaturization examples of TiLV polymerase PB1 subunit. Corresponding structures of TiLV PB1 (46-73) and FluPol PB1 (84-123) are coloured in light blue and dark blue. **c.** Absence of β-ribbon in TiLV PB1 subunit. Corresponding structures of TiLV PB1 (107-137) and FluPol PB1 β-ribbon (162-222) are coloured as in **b**. Flexible residues are shown as dotted line. **d.** TiLV PB1 subunit does not have a priming loop. TiLV and FluPol PB1 thumb domains are respectively coloured in light and dark aquamarine. FluPol priming loop is coloured in red. Flexible residues are represented as a dotted line. **e.** TiLV PB1 active site. RdRp motifs A, B, C, D, E and F are respectively coloured in dark turquoise, magenta, dark grey, light green, light grey and blue. Key residues are displayed and the active site aspartates are circled. In the vRNA initiation state, CTP (dark grey) is in +1 active site position. Magnesium ions (Mg^2+^) are shown as green spheres. The NTP entry, template entry and template/product exit channels are indicated with arrows.

Despite being minimal, the TiLV PB1 subunit contains all the conserved RdRp functional motifs (Fig. 3e). Motifs A (208-223) and C (285-295) are implicated in divalent metal ion coordination through catalytic aspartic acids D213, D289 and D290. Motif B (263-278) is characterized by an *Articulavirales*-conserved methionine rich loop (262-GG<u>M</u>L<u>MM</u>FN). It is implicated in stabilization of the incoming NTP at the +1 position (M266), in translocation and in discrimination of NTPs from dNTPs (N270). Motif D (311-323) contributes to a positively charged entry tunnel for NTPs via lysine residues K318 and K319, in concert with K216 (motif A) and K151 and R155 (motif F). Motif E (326-335) stabilizes motif C through hydrophobic interactions (L327, I329). Motif F (138-176) is also implicated in template stabilization (I157, R165).

### TiLV PB2 subunit

TiLV PB2 subunit (457 residues) is 40 % smaller than FluPol PB2 (760 residues), but similarly divided into the PB2 N-terminal domain (PB2-N, 1-145) and an array of PB2-C domains. These include the split mid-link (mid: 146-171, link: 245-280), CBD-like (172-244), 627-like (281-380) and NLS (381-457) domains (Fig. 4a). TiLV PB2-N contains a minimal two α-helical lid domain (77-117, 41 residues) compared to the larger FluPol one (80 residues), which is implicated in template-product duplex separation during elongation^23^ (Fig. 4b).

**Figure 4.**
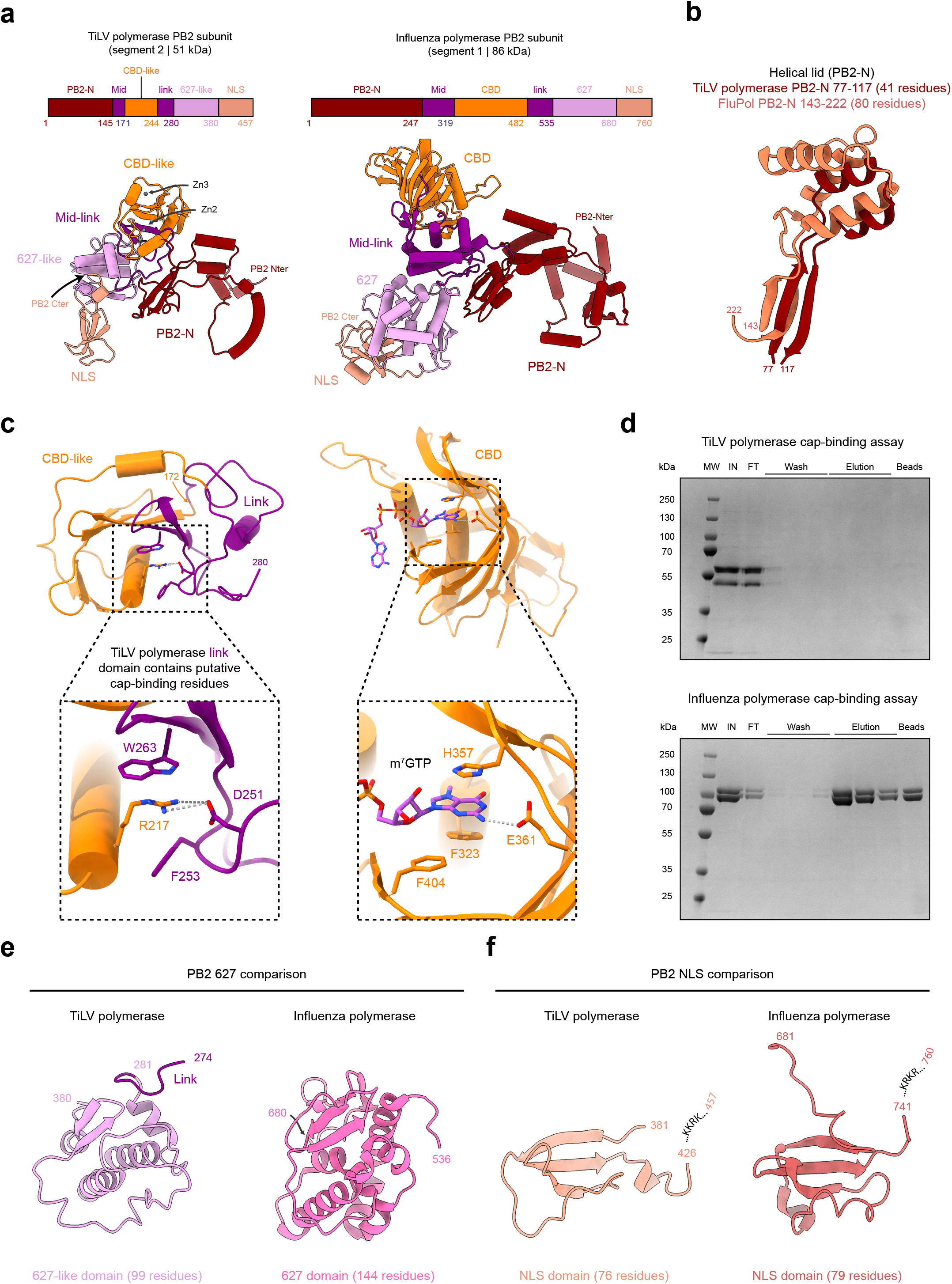
TiLV polymerase PB2 subunit in transcriptase conformation. **a.** Schematic and cartoon representation of TiLV polymerase and FluPol PB2 subunits in transcriptase conformation. Domains are coloured as in Fig. 1c. **b.** Miniaturization example of TiLV PB2-N lid. TiLV polymerase lid (77-117), coloured in dark red, is superposed with FluPol lid (143-222), coloured in orange. **c.** Comparison of TiLV PB2 CBD-like and FluPol PB2 CBD. **Top:** Overall structures of TiLV PB2 CBD-like and link domains (245-280) and FluPol PB2 CBD. **Bottom:** Close-up views of each respective cap-binding site. TiLV polymerase CBD-like and link domains enclose putative cap-binding residues. FluPol CBD is bound to m^7^GTP (light blue). **d.** *In vitro* TiLV polymerase cap-binding assay. **Top:** SDS-PAGE analysis of TiLV polymerase cap-binding capacity using immobilized γ-aminophenyl-m^7^GTP (C10-spacer) beads. The molecular ladder (MW) is shown on the left of the gel. “IN” corresponds to the input. “FT” corresponds to the flow through. **Bottom:** SDS-PAGE analysis of A/H7N9 FluPol cap-binding capacity using immobilized γ-aminophenyl-m^7^GTP (C10-spacer) beads annotated as above. **e.** Comparison of TiLV PB2 627-like and FluPol PB2 627 domains. **f.** Comparison of TiLV PB2 NLS and FluPol PB2 NLS domains. The respective NLS sequence is indicated. In both cases, the NLS C-terminal is not visible due to flexibility.

The minimized CBD-like domain is inserted into the mid-link domain and, unusually, both domains contain putative cap-binding residues. F253 and W263 from the TiLV link domain form a potential aromatic sandwich comparable to PB2/H357 and F404 in FluPol, between which stacks the m^7^G cap, and TiLV D251 is equivalent to FluPol PB2/E361, which ensures guanine specificity (Fig. 4c). However, in TiLV, R217 is sandwiched between F253 and W263 and makes a salt bridge with D251. The auto-inhibition of potential cap-binding by R217 in TiLV is reminiscent of Thogoto virus (THOV) polymerase^21^ and severe fever with thrombocytopenia syndrome virus (SFTSV, also called Dabie bandavirus) L protein^24, 25^. For THOV CBD-like domain, PB2/R344 is sandwiched between an aromatic and an arginine and for SFTSV-L CBD, R1843 is between a pair of aromatic residues (Extended Data Fig. 3). THOV CBD-like domain appears to be non-functional^21^, but in the case of SFTSV, it has been shown structurally that R1843 can swing out of the way to allow cap-binding during transcription initiation (Extended Data Fig. 3). To test whether TiLV polymerase can bind cap, we used immobilized m^7^GTP beads. We found that vRNA promoter bound TiLV polymerase is not able to bind the beads, in conditions where FluPol can (Fig. 4d). It remains to determine whether cap binding is possible under different conditions.

The TiLV 627-like domain (281-380, 99 residues) is substantially smaller than the equivalent FluPol domain (144 residues) but has the same topology (Fig. 4e), whereas the TiLV NLS domain (381-457, 76 residues) is of comparable size to the FluPol NLS domain (79 residues) (Fig. 4f). The folded part of the TiLV NLS domain is immediately followed by a putative NLS motif 427-KKRK, similar to FluPol 740-KRKR. In all TiLV polymerase cryo-EM maps, PB2 C-terminal residues 425-457 remain unseen, presumably due to their flexibility. In FluPol, a stable PB1-PB2 inter-subunit interaction is formed by a helical bundle involving the C-terminus of PB1 and the N-terminus of PB2, reinforced in the transcriptase conformation by a helix from the endonuclease domain. In TiLV, similar interactions are observed although much reduced in scale (not shown). Perhaps to compensate, in TiLV there is a unique extra C-terminal extension to PB1 (494-515) that wraps around and reinforces the small PB2 helical lid.

### TiLV polymerase binding to the vRNA and cRNA promoters

As for other sNSVs polymerases^10–14^, TiLV polymerase can bind both the vRNA and cRNA promoter in two distinct modes (Fig. 5). In mode A, the 3’ end is directed into the polymerase active site for initiation of RNA synthesis (Fig. 5a; Extended Data Fig. 4), whereas in mode B, the 3’ end is docked on the surface of the TiLV polymerase core, in the so-called “secondary site” (Fig. 5e; Extended Data Fig. 5).

**Figure 5.**
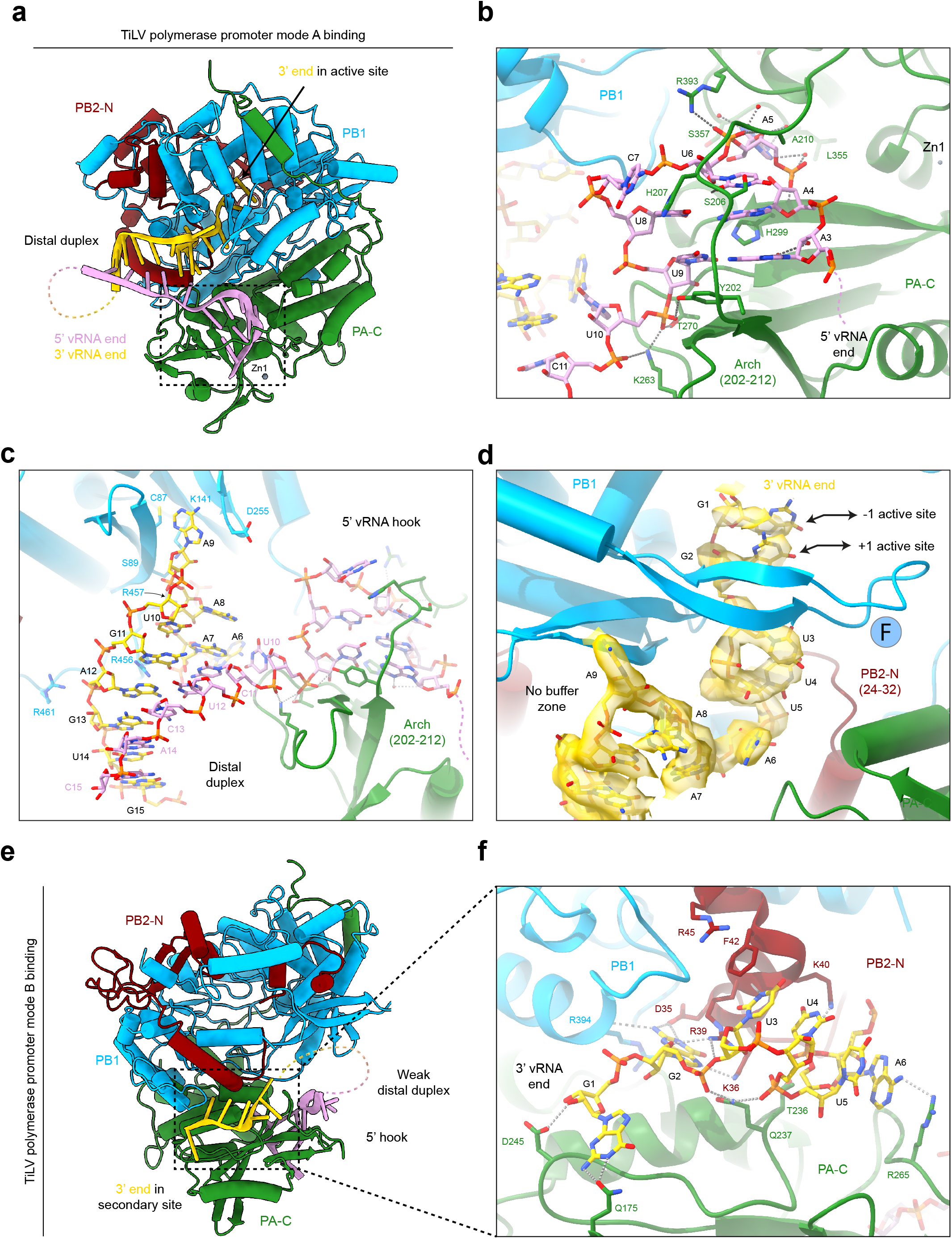
TiLV polymerase binding to the vRNA promoter. **a.** Cartoon representation of TiLV polymerase bound to the 40-mer vRNA loop in promoter mode A. PA is coloured in green, PB1 in light blue and PB2-N in dark red. The 5’ vRNA end is in pink, the 3’ vRNA end in gold. The 3’ end is directed towards the polymerase active site. The 5’/3’ ends form a distal duplex with the flexible linker nucleotides shown as a dotted line. **b.** Close-up view of the bound 5’ vRNA hook. TiLV polymerase subunits are coloured as in The two first 5’ end nucleotides are flexible and represented as a dotted line. The PA arch (202-212) stabilizing the 5’ hook structure is indicated. Water molecules are red spheres and hydrogen bonds grey dotted lines. Key interacting residues are shown as sticks. **c.** Close-up view of the distal duplex formed between the 5’ and 3’ vRNA ends with annotation as in **b**. 3’ end nucleotides are numbered from the 3’ end, with nucleotides 1-5 omitted for clarity. **d.** Close-up view of the vRNA 3’ end entering the TiLV polymerase active site in pre-initiation state mode A, with annotation as in **c**. The RdRp motif F β-hairpin is shown, as well as the -1/+1 active site positions occupied respectively by G1 and G2. **e.** Cartoon representation of TiLV polymerase core bound to the 40-mer vRNA loop in promoter mode B with annotation as in **a**. The 3’ end is bound to the polymerase surface in the so-called “secondary site”. The 5’/3’ ends form a weak distal duplex in the closed core conformation. **f.** Close-up view of the 3’ vRNA end bound in the secondary site with key interacting residues shown as sticks. Annotation and colours as in **b** and **c**.

In both modes, the vRNA 5’ end nts 3-AAAUCUU-9 form a compact stem-loop structure (hook) that binds in a pocket covered by the PA arch (202-212) (Fig. 5b). A3 and A4 respectively base pair with U9 and U8, while A5 to C7 form a loop with U6 stacking on A4 (Fig. 5b; Extended Data Fig. 4a, 5a). The two first 5’ end nucleotides have no density and are presumably flexible. The 5’ hook interacts with the polymerase through backbone interactions and base stacking (e.g. PA/H207 on C7 and PA/Y202 under the A3-U9 base pair). Only splayed out A5 and U6 are base-specifically recognized by PA/L355 and S357 and PA/S206 and H207 respectively (Fig. 5b; Extended Data Fig. 4a). The 5’ hook position and shape is similar to that observed in other sNSV viruses but is unusually small, comprising only 7 nts compared with 10 for FluPol, LACV-L/SFTSV-L, LASV-L and up to 12 for HTNV-L (Extended Data Fig. 6). In the vRNA mode A pre-initiation state, a strong 5-mer distal duplex is formed by 5’ end nucleotides C11 to C15 base-pairing with 3’ end nts G11 to G15 (Fig. 5c). U10 connects the hook to the distal duplex, with a sharp bend being induced by the binding of K263 to the phosphates of both U10 and C11 (Fig. 5bc, Extended Data Fig. 4a). The 3’ end enters the active site in a highly structured way (Fig. 5cd, Extended Data Fig. 4a) with A9 and U5 being splayed base-specific recognition pockets. A8 stacks on A7 and U4 and U3 are stacked on each other with U3 being base-specifically recognized. Finally, G1 and G2 are stacked on each other at the -1 and +1 positions respectively in the polymerase active site (Fig. 5d, Extended Data Fig. 4a).

There are six differences in the vRNA and cRNA promoter sequence for segment 9, which however do not significantly affect the overall mode A or mode B promoter structure (Extended Data Fig. 4a, 5a). Notably, all base-specifically recognized nucleotides (5’ A5 and U6 and 3’ U3, U5 and A9 as well as G2 in mode B, see below) are absolutely conserved in the vRNA and cRNAs of all segments. Only the substitutions from 5’ U10 and 3’ A6 (vRNA) to 5’ A10 and 3’ G7 (vRNA) cause a locally different nucleotide packing against the distal duplex in mode A, other base substitutions being accommodated without perturbation. The trajectory of the 3’ end into the active site is identical for vRNA and cRNA, except that for cRNA, C1 and G2 are at the -1 and +1 positions respectively, rather than G1 and G2 for vRNA. This suggests that for both cRNA and vRNA, replication is initiated terminally. Whereas other sNSVs polymerases have a ‘buffer zone’ that allows the template to extend and then retract^26^, while maintaining the distal duplex during prime-and-realign, the extensive and tight binding of the 3’ end suggests this might not be the case in TiLV.

In the mode B pre-initiation state, the 5’ hook binds as in mode A, but 3’ end nts 1-GGUUUA in vRNA are sequestered in the secondary site at the interface of PA-C, PB1 and PB2-N on the surface of TiLV core (Fig. 5e; Extended Data Fig. 5). G1 is stabilized by PA/Q175 and D245, while G2 is base-specifically bound in a pocket, stacking between PA/M229, R233 and PB2/R39, with PB2/D35 and K36 contacting N2 and N7 respectively and the carbonyl of PB1/R394 contacting N2. U3 is sandwiched between PB2/F42 on one side and U4 on the opposite side, with PA/Q237 bridging both G2 and U4 phosphates (Fig. 5f; Extended Data Fig. 5a). Extensive 3D classification allowed two distinct populations of mode B vRNA particles to be isolated (Supplementary Information 4), either with a closed core and ill-defined distal duplex, due to disorder of 3’ end residues A7 to G15 (Extended Data Fig. 5a, b), or with an open core and well-resolved distal duplex (Extended Data Fig. 5a, c). For cRNA, only the closed core conformation without distal duplex is observed (Extended Data Fig. 5d, Supplementary Information 2). In the open conformation, vRNA 3’ end A6 is stabilized in a pocket formed by PA/T236, T267, T270 and H272 residues. A7 interacts with PA/R265 and stacks with A8. A9 is not visible but its flexibility allows U10 to flip out and stack with PA/R265, the latter being stabilized by R194 (Extended Data Fig. 5a, c). The distal duplex is formed, as in mode A, between the 3’ vRNA nucleotides G11 to G15 base-pairing with the 5’ vRNA nucleotides C15 to C11 (Extended Data Fig. 5a, c). It is notable that the mode B duplex is at a significantly different orientation to that observed in mode A (Extended Data Fig. 5e). The two mode B conformations may represent intermediate states in the recycling from mode B to mode A at the end of RNA synthesis to allow a new round of RNA synthesis, as proposed for FluPol^27^.

### TiLV polymerase activity: from pre-initiation to initiation

We next investigated whether the polymerase is able to perform *de novo* replication-like or primed transcription-like activity *in vitro*. Based on the pre-initiation mode A structures, which shows 3’ terminal nucleotides G1-G2 (vRNA) or C1-G2 (cRNA) respectively in positions -1/+1 of the active site (Fig. 6a), we hypothesized that *de novo* replication would likely initiate by the synthesis of the dinucleotide pppCpC or pppGpC, from CTP+CTP or GTP+CTP for vRNA or cRNA, respectively.

**Figure 6.**
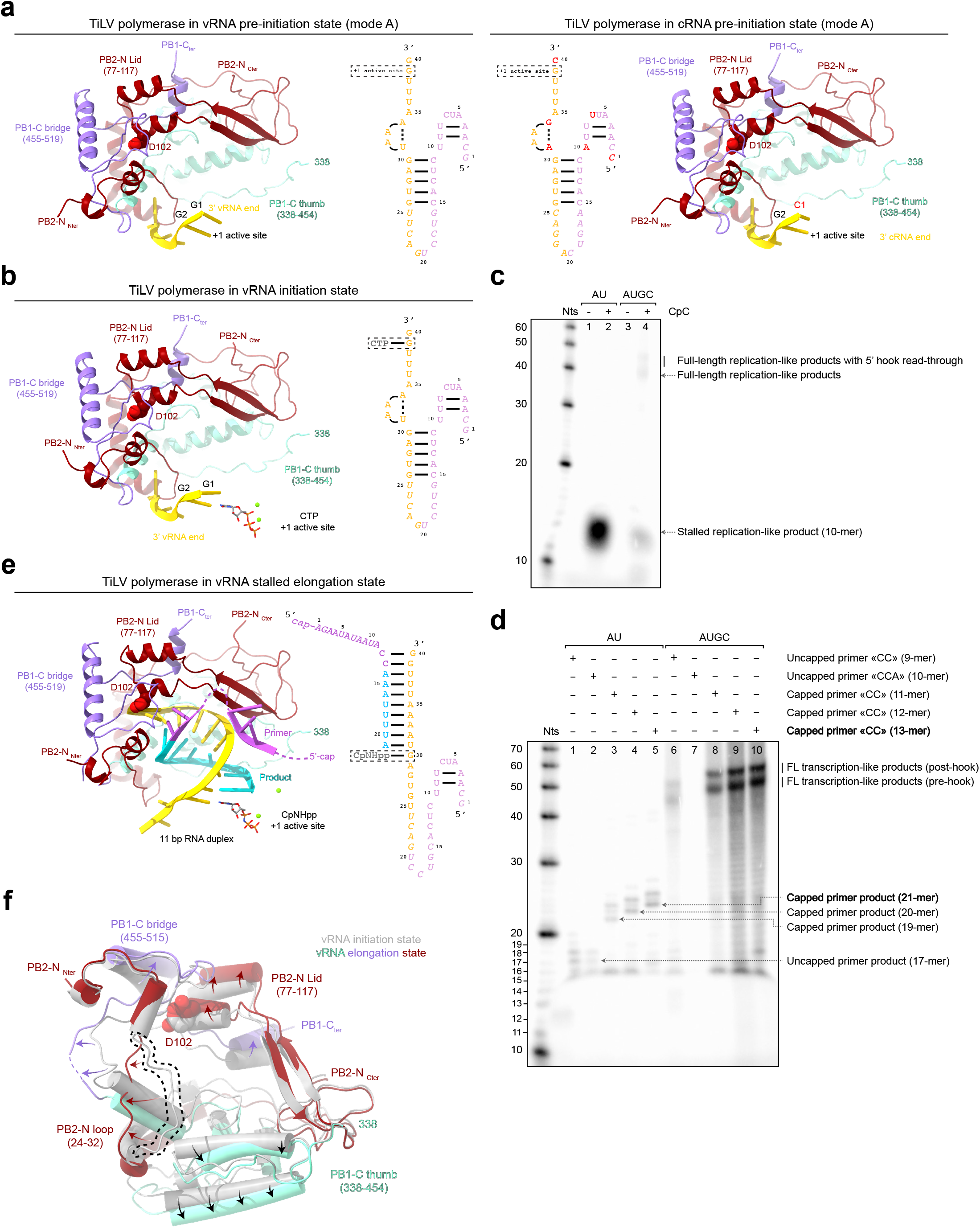
TiLV polymerase RNA synthesis activity: from-pre-initiation to elongation. **a.** TiLV polymerase in vRNA and cRNA pre-initiation state mode A. **Left:** Close-up view of the 3’ vRNA end in the TiLV polymerase active site. The 3’ end is coloured in gold and nucleotides are numbered from the 3’ to the 5’, with G1 and G2 being in respectively the -1/+1 active site positions. PB1-C bridge domain is coloured in purple, PB1-C thumb domain in aquamarine and PB2-N in dark red. PB2-N/D102 from the lid domain is shown as spheres. Schematic of the overall vRNA conformation is shown on the right with the +1 active site position indicated with a dotted rectangle. Flexible nucleotides are in italic. **Right:** Close-up view of the 3’ cRNA end in the TiLV polymerase active site, annotated as in **a,** except that C1 and G2 are respectively in the -1/+1 active site position. Schematic of the overall cRNA conformation is shown on the right with the +1 active site position indicated with a dotted rectangle. Nucleotide differences between the vRNA and the cRNA are coloured in red. **b.** TiLV polymerase in the vRNA initiation state with incoming CTP aligning with G2, annotated as in **a**. Magnesium ions (Mg^2+^) are shown as green spheres. Schematic of the overall RNA conformation is shown on the right with the +1 active site position and the CTP indicated with a dotted rectangle. Flexible nucleotides are in italic. **c.** Urea-PAGE analysis of *in vitro* TiLV polymerase replication-like activity using the vRNA loop template and ATP and UTP (AU), without (**lane 1**) or with (**lane 2**) CpC dinucleotide. The expected stalled replication-like product (10-mer) is indicated on the right. Replication-like activity assay using all NTPs (AUGC), without (**lane 3**) or with (**lane 4**) CpC dinucleotide. Expected full-length replication-like products are indicated on the right. The decade marker (Nts) is shown on the left. Uncropped gel is provided as source data file. **d.** Urea-PAGE analysis of *in vitro* TiLV polymerase transcription-like activity using ATP and UTP (AU), uncapped primers ending by …CC-3’ (**lane 1**) or …CCA-3’ (**lane 2**), capped primers ending by …CC-3’ of different lengths (11-mer, **lane 3**; 12-mer, **lane 4**; 13-mer, **lane 5**). Expected stalled transcription-like products are indicated on the right. Transcription-like activity assay using all NTPs (AUGC), uncapped primers ending by …CC-3’ (**lane 6**) or …CCA-3’ (**lane 7**), capped primers ending by …CC-3’ of different lengths (11-mer, **lane 8**; 12-mer, **lane 9**; 13-mer, **lane 10**). Expected full-length transcription-like products are indicated on the right. The decade marker (Nts) is shown on the left. Uncropped gel is provided as source data file. **e.** TiLV polymerase in the stalled elongation state showing the internal 11 base pair (bp) product-template RNA duplex. Domains and residues are annotated as in **a**. The 3’ vRNA end is coloured in gold, not yet pushed back by PB2-N D102 lid residue, shown as spheres. The capped primer ending by …CC-3’ (13-mer) is coloured in magenta and the incorporated nucleotides are in cyan. Flexible nucleotides are shown as dotted lines. A CpNHpp is in the +1 active site position. Magnesium ions (Mg^2+^) are shown as green spheres. Schematic of the overall RNA conformation is shown on the right with the +1 active site position and the CpNHpp indicated with a dotted rectangle. Flexible nucleotides are in italic. **f.** TiLV polymerase movements upon product elongation. The vRNA initiation state (grey) was superposed with the elongation state (colours) taking PB1 subunit as reference. The RNA is not displayed for clarity. Only significant movements of elements of PB1-C and PB2-N are shown, and highlighted with arrows. The PB2-N loop (24-32) in the vRNA initiation state, which has to displace to allow elongation, is surrounded by a dotted line.

To test this hypothesis, we incubated TiLV polymerase with the vRNA loop, CTP, Mg^2+^ and froze the reaction on grids after 1h. We obtained two cryo-EM structures of TiLV polymerase in the vRNA initiation-like state, one with the complete polymerase (2.73 Å resolution) and one with the polymerase core only (2.4 Å resolution, enabling visualization of many water molecules) (Extended Data Fig. 4d, Supplementary Information 1). The initiation-like state is very similar to the pre-initiation mode A structure, except that a single CTP with two magnesium ions is bound at the +1 active site position, base pairing with template nucleotide G2. Binding of CTP induces only small perturbations in the tilt of the template nucleotides G1 and G2, in the motif C loop, enabling co-ordination of the active site aspartates with the two catalytic magnesium ions and in some motif F residues (PB1/K151 and R155) that co-ordinate the triphosphate (Fig. 6b, Extended Data Fig. 7c). Unlike FluPol^23^, there is no major switch in the conformation of the motif B loop backbone, although M266 side chain does flip under the CTP base. The absence of the dinucleotide 5’-pppCpC hybridized to the 3’ vRNA end suggests that it does not form under these conditions or is too weakly bound, bearing in mind that there is no priming loop that might stabilize the pre-catalytic complex. Nevertheless, the template and CTP positions are consistent with TiLV polymerase replication initiation being terminal (Fig. 6a, b; Extended Data Fig. 7).

### TiLV polymerase activity: from initiation to elongation

We next established *in vitro* assays to characterize *de novo* and primed RNA synthesis activity by TiLV polymerase (Fig. 6cd). Using the vRNA loop and all NTPs, we failed to observe *de novo* reaction products (Fig. 6c, lane 3). Given that we did not observe the 5’-pppCpC dinucleotide product in the initiation state structural analysis, this suggests that *de novo* replication is limited by the initiation step. We therefore used the dinucleotide CpC as a primer to hybridize to the vRNA template 3’ end …GG-3’, equivalent to the use of ApG in studies of FluPol RNA synthesis. CpC together with ATP and UTP only, resulted in strong production of the expected 10-mer product, stalled through lack of CTP to match G11 of the template (Fig. 6c, lane 2). Using CpC with all NTPs yields two weak bands of around 40 nts that likely correspond to full-length products, with or without read-through of the 5’ hook (Fig. 6c, lane 4).

As we do not know whether cap-binding occurs nor the preferred length of a putative capped primer, to investigate transcription-like behavior we used capped (11-, 12-, 13-mers; all ending with …CC-3’) and uncapped (9-mer, ending with …CC-3’; 10-mer, ending by …CCA-3’) primers to initiate RNA synthesis, with either ATP and UTP only or all NTPs (Fig. 6d). Using both uncapped primers in combination with ATP, UTP, and Mg^2+^, we observe the synthesis of the expected 17-mer product due to stalling at G11, but also slightly longer products that could result from read-through due to misincorporation, leading to stalling at G13 or G15 of the template (Fig. 6d, lane 1-2). Similar results are obtained with the three capped primers (Fig. 6d, lane 3-5). With all NTPs, two long products, differing by about 9 nts, are seen with all primers ending in …CC-3’ (Fig. 6d, lane 6,8-10), consistent with termination pre- or post-hook. Interestingly, reactions with the uncapped …CCA-3’ primer show lower intensity (with ATP and UTP) or no product (with all NTPs) compared to the uncapped CC-3’ primer (Fig. 6d, lane 1-2, 6-7). This suggests that U3 of the vRNA template is poorly accessible for hybridization with a primer, consistent with its specific recognition in a pocket distinct from the active site (Fig. 5d) and highlighting the inflexible positioning of the template 3’ end and the absence of a template buffer zone. This is not the case for FluPol, for which capped primers ending by …AG-3’ or …AGC-3’ are equally reactive since the template is advanced by one position into the active site^23, 27^. While we do not detect any cap-binding *in vitro*, reactions with the 12- and 13-mer capped primers show stronger product formation, which could indicate the optimal primer length used by TiLV polymerase for transcription initiation. These preliminary results give the first insights into TiLV polymerase replication and transcription-like activities, but a more detailed analysis is required to understand for instance the requirements for efficient replication initiation and the mechanism of termination, with or without polyadenylation.

To structurally characterize the early elongation state, TiLV polymerase was incubated with the vRNA loop, a capped 13-mer primer ending in …CC-3’, ATP, UTP, Mg^2+^ and CpNHpp. This leads to elongation of the primer by eight nucleotides to become a 21-mer, before stalling at nucleotide G11 of the template (Fig. 6e). The reaction products were applied to EM grids after 4 h, frozen and two distinct elongation structures of TiLV polymerase were determined from cryo-EM data at respectively 2.96 and 2.42 Å resolution (Table 1, Extended DataTable 1, Extended Data Fig. 8ab, Supplementary Information 4). Both show the full-length TiLV polymerase with CpNHpp bound at the +1 position and a product-template duplex filling the active site cavity (Fig. 6e). Unambiguous RNA sequence identification, possible because of the high resolution of the cryo-EM density, confirms that the observed elongation state is consistent with the biochemistry (Extended Data Fig. 8ac). The higher resolution structure differs in having an additional 3’ end bound in mode B forming a distal duplex with the 5’ end (see below). Unusually, the product-template duplex comprises 11 base pairs (+1 to -10) (Fig. 6e) where generally for viral polymerases a maximum of 10 base pairs is observed. This likely arises from the fact that the PB2 lid domain has not yet managed to enforce strand separation. Details of the interactions of the RNA within the active site cavity are shown in Extended Data Fig. 8d. The CpNHpp is bound is a non-catalytic configuration with only one magnesium ion. Although a capped primer was used to initiate the elongation reaction, the cap itself is not observed and the putative cap-binding site remains autoinhibited by PB2/R217. However, density tentatively assigned to primer nucleotides 4-6 is observed in the presumed product exit channel at the interface of PB1 and PB2, with A6 stacking on PB2/H305 (Extended Data Fig. 8abd). In the absence of a priming loop, TiLV polymerase template exit tunnel always remains open, different from FluPol (*Articulavirales*) or LACV-L (*Bunyavirales*) in which the RNA duplex extension induces the extrusion of the priming loop or the template exit plug, respectively^23, 26^ (Extended Data Fig. 9). Overall, compared to the initiation state, the PB1 thumb (338-454) and bridge (455-515) and PB2-N, including the loop 24-32 and the lid (77-117), rotate relative to PA and the N-terminal region of PB1, expanding the active site cavity to accommodate the product-template duplex (Fig. 6f). The higher resolution structure with the additionally bound 3’ ends mimics a later elongation state in which the outgoing template would have translocated into the 3’ end secondary site, suggesting that this template trajectory, facilitating template recycling for further rounds of transcription/replication^27^, is a conserved feature of all sNSV polymerases (Extended Data Fig. 9).

### Conformational flexibility of TiLV polymerase

For each major functional state described above, additional cryo-EM structures were determined that exhibit different positons of some of the flexible linked domains (Table 1). Of particular interest is the observation of a replicase-like configuration of TiLV polymerase in which the ENDO-like and PB2-C domains are rearranged as previously described for FluPol (Fig. 1d, Extended Data Fig. 10ab)^17–20^. This structure occurred as a subpopulation in the elongation state sample (Supplementary Information 4) and has the vRNA promoter bound in mode A and initiating CpNHpp is at the +1 position base pairing with template nucleotide G2. The replicase conformation is so-called as it occurs in an asymmetric dimer with the encapsidase in the presumed influenza replication complex^20^. In the transcriptase conformation, the PA endonuclease faces the PB2 cap-binding domain, in a configuration compatible with cap-snatching (Extended Data Fig. 10c), with the PB2 627 and PB2 NLS domains contacting each other and interacting with PB1 (Extended Data Fig. 10d). To switch from the transcriptase conformation to the replicase conformation, the TiLV ENDO-like domain rotates by ∼160 degrees and the PB2 NLS domain separates from the 627 domain (which remains flexible) to pack against the ENDO-like domain, interacting through hydrophobic and electrostatic interactions (Extended Data Fig. 10c). Concomitantly, the PB2 CBD-like and midlink domains rotate by ∼180 degrees to pack against TiLV PB2-N and PB1 (Extended Data Fig. 10c). The C-terminal extension to the folded part of the TiLV NLS domain (i.e. beyond residue 420), which actually bears the putative NLS, is not observed in any of the TiLV structures. In contrast, in the FluPol replicase conformation, the corresponding region forms an alpha helix interacting with the endonuclease domain (Fig. 1e)^17–19^. In some of the TiLV cryo-EM structures the full replicase configuration is not observed, only the rotated endonuclease with or without associated NLS domain (Table 1), highlighting the flexible nature of these peripheral domains. Finally, we note that for the transcriptase conformation, there is a systematic difference between complete polymerase structures (i.e. including ENDO-like and PB2-C domains) and those of the core only. Packing of the 627-like domain against PB1 induces a conformational change that propagates towards the active site (Extended Data Fig. 10e), although the significance of this is unclear.

## Discussion

In this study, we structurally and functionally characterized the heterotrimeric TiLV polymerase in complex with both the vRNA and cRNA promoters. We thus report the first complete structure of a polymerase from the *Amnoonviridae*, an evolutionary distant viral family from the O*rthomyxoviridae* (which encompasses Influenza, Thogoto, Quaranja and Isaviruses), both within the *Articulavirales* order. Recently, metagenomics^28^ has revealed a number of other *Amnoonviridae* species, mainly from fish, including Flavolineata, Namensis, Asotus1, Asotus2, Przewalskii and Stewartii viruses^29, 30^. Sequence alignments of the putative polymerase segments (full-length sequences only, Supplementary Data 5-8), shows, as expected, reasonable conservation of the PB1 subunit, but significant divergence in the PB2 and PA segments, indicating sparse sampling of this family to date^30^. For PB1, TiLV, Flavolineata, Namensis are the shortest (∼520 aa) and lack ∼24 residue insertions in both the fingers and thumb compared to Asotus1, Asotus2, Przewalskii and Stewartii (∼ 567 aa) (highlighted in blue, Supplementary Data 5). For PB2, TiLV and Namensis have significant homology, with zinc binding residues and most but not all putative cap-binding or inhibiting residues conserved (Supplementary Data 6), whereas the other four (Asotus1, Asotus2, Przewalskii and Stewartii) form a divergent clade, which the Zn2 site is conserved but not Zn3 and the putative NLS sequence is only apparent in TiLV (Supplementary Data 7). For PA, there is very low overall homology between the three full-length sequences (TiLV, Asotus1, Flavolineata), with the Zn1 binding residues being conserved but not the putative endonuclease ‘active’ site residues (Supplementary Data 8). Most recently, metagenomics of corals in addition to transcriptome mining has led to identification of a proposed new family, *Cnidenomoviridae*, within the *Articulavirales* order, which would now include at least four families: *Orthomyxoviridae*, *Amnoonviridae*, *Quaranjaviridae*, and *Cnidenomoviridae* (Petrone et al, https://www.biorxiv.org/content/10.1101/2023.02.15.528772v1.full.pdf). Future analysis of additional sequences will undoubtedly shed more light on the evolution of *Articulavirales* polymerases.

To date, TiLV polymerase is the smallest known viral polymerase from the sNSV group with a molecular weight of only 155 kDa, around 60 % the size of FluPol or *Bunyavirales* L proteins. The putative viral nucleoprotein (segment 4) is also considerably smaller than its influenza counterpart^31^. Remarkably, the small polymerase size is achieved by an overall pruning of each subunit whilst maintaining all the functional domains typically found in orthomyxo-like viral polymerases. This is achieved in a variety of ways as described above (Figs. 2-4). A small polymerase size (and small size of gene segments in general, at least for TiLV) seems to be a feature of the *Amnoonviridae*. Given that bunya- and orthomyxo-viruses have similar sized, large polymerases, yet are probably more distantly related, it suggests that some evolutionary selective pressure may have led to a general gene size reduction in *Amnoonviridae*, perhaps the need to package ten distinct proteins/segments in the case of TiLV. Nevertheless, the minimal TiLV heterotrimeric complex can take up active transcriptase- or replicase-like conformations, equivalent to those found for FluPol. However, we find that the TiLV ENDO-like domain does not have an obvious metal-binding active site and is unable to cleave RNA *in vitro* (Fig. 3). Similarly, the TiLV CBD-like domain does not bind m^7^GTP *in vitro*, which could be explained by consistent autoinhibited by PB2/R217 (Fig. 4). However, these anomalies are shared with some other sNSV viral polymerases (Extended Data Fig. 2, 3). For example, the isolated Thogoto and Dhori virus PA-ENDO like domains do not exhibit any catalytic residues and are inactive *in vitro* and the Thogoto PB2 CBD-like domain is autoinhibited and does not bind cap *in vitro*^21^. Similarly, the Lassa virus CBD-like domain does not bind cap *in vitro*^32, 33^. On the other hand, the SFTSV-L CBD undergoes a conformational change that releases auto-inhibition to enable 5’-cap binding and cap-dependent transcription^25^ (Extended Data Fig. 3). Whether or not, TILV polymerase performs cap-snatching and if so, how, remains to be elucidated.

Like all sNSV polymerases, TiLV polymerase binds the 5’ end promoter region of vRNA or cRNA as a hook, although it is exceptionally small (nucleotides 3-9) with the first two 5’ end nucleotides overhanging. The 3’ end (vRNA and cRNA) can bind in the RdRp active site (mode A) or in the secondary site the surface of TiLV polymerase core (mode B) (Fig. 5). This suggests that the trajectory of the translocating template and the mechanism of template recycling is as first described for FluPol^27^, but common also to *Bunyavirales*. However, there is no conserved oligo-uridine sequence close to the 5’ end (Fig. 1b) that could act as a polyadenylation signal, as in influenza virus. The mechanisms of transcription termination and potential poly-adenylation therefore still need to be elucidated. Finally, it is interesting to note that there is no Watson-Crick complementarity between the first position of the 5’ and 3’ ends (G:G, C:C for vRNA and cRNA, respectively, Fig. 1b). Perhaps this allows escape from anti-viral innate immune receptors such as RIG-I that depend on recognition of a canonical blunt-end 5’ppp-dsRNA^34^.

*In vitro* RNA synthesis assays and structural data on initiation and elongation states show that promoter-bound TiLV polymerase, despite its reduced size, is able to perform both replication-like and transcription-like activity in canonical fashion (Fig. 6). However, some crucial structural elements are missing when compared to FluPol. The absence of a priming loop and the fact that both 3’ vRNA and cRNA ends align in the active site consistent with terminal initiation suggests that the TiLV replication initiation mechanism may be simpler than for other sNSV polymerases, which rely heavily on prime-and-realign^14^. For FluPol, vRNA to cRNA initiation is terminal (requiring the priming loop), while cRNA to vRNA synthesis initiates internally, with the use of a potential trans-activating apo-polymerase for template realignment^19^.

Overall, our results provide the first structural and functional data on a prototypic viral polymerase from the *Amnoonviridae* family that show functionality is not compromised despite an unprecedented reduction in size. Unfortunately, the current lack of reverse genetics or a minireplicon system for TiLV limits the extent to which viral RNA synthesis mechanisms can be studied and tested in cells. However, our highlighting of conserved and idiosyncratic features compared to classical orthomyxoviruses will contribute to establishing such systems, pave the way for a better understanding of TiLV replication and perhaps help combat this emerging pathogen of tilapia fish, an important human food resource.

## Methods

### Cloning, expression and purification of the TiLV polymerase heterotrimeric complex

The 10 TiLV open reading frames, codon optimized for insect cells expression, were ordered at Genscript and sub cloned into multiple pFastBac Dual vectors using EcoRI and SpeI (pFastBac Dual SEGX, with X being the segment number). Plasmids containing each single segment were PCR amplified using primers sequences 5’-CGCTGAGCAATAACTATCATAACCCCTAGGAGATCCGAACCAGATAAGTG-3’ and 5’-GGTTATGATAGTTATTGCTCAGCGCTCAAGCAGTGATCAGATCCAGACATG-3’, re-circularized by Gibson cloning thus creating 10 pFastBac plasmids compatible with the biGBac cloning system^35^ Single cassettes encoding for the 10 TiLV segments were amplified using the biGBAc standard primers and assembled by Gibson assembly into psBIG1a and psBIG1b. psBIG1a and psBIG1b were linearized with Swa1, resulting in psBIG1a TiLV 1-3-5-7-9 and psBIG1b TiLV 2-4-6-8-10. Finally, psBIG1a TiLV 1-3-5-7-9 and psBIG1b TiLV 2-4-6-8-10 were digested using Pme1 and fragments encoding the 10 TiLV proteins were cloned by Gibson assembly into psBIG2ab, linearized using Pme1, to finally give psBIG2ab TiLV-10SEG.

The construct encoding for segment 1, segment 2 (with a Tobacco Etch Virus (TEV) protease cleavage site followed by a 10x-histidine tag on the C-terminus) and segment 3 proteins was obtained as following:

1. A pFastBac Dual compatible with the biGBac system was digested with EcoR1 and Spe1 as well as the coding sequence of segment 1, PCR amplified using primers 5’-ATACCGTCCCACCATCGGGC-3’ and 5’-CGCGACTAGTTTAGCATCCAGCAGTAGGCTGGAC-3’. Insert and backbone were ligated using T4 DNA ligase (NEB) following manufacturer recommendations giving the pFastBac Dual TiLV-SEG1-noHis.
2. Segment 2, PCR amplified using primers 5’-ATACCGTCCCACCATCGGGC-3’ and 5’-GCGACTAGTTTAGTGGTGGTGGTGATGATGGTGATGATGGTGTCCCTGGAAGTACAGGTTTTCTGAGCCAGAACCCTGGTTCAGGTCCATGTTTTCAGC-3’, was digested using EcoR1 and Spe1 for subsequent ligation into a pFastBac Dual opened with EcoR1 and Spe1 using the T4 DNA ligase (NEB) following manufacturer recommendations, giving the pFastBac Dual TiLV-SEG2-TEV-His.
3. Finally, cassettes encoding for TiLV-SEG1-noHis, TiLV-SEG2-TEV-His and TiLV-SEG3 were amplified using standard biGBac primers and assembled by Gibson cloning into psBIG1a, opened by Swa1, and obtain psBIG1a TiLV-SEG1-noHis_SEG2-TEV-His_SEG3.

EMBacY bacmids^36^ containing the TiLV polymerase expression plasmid were generated and used for insect cells expression. For large-scale expression, *Trichoplusia ni* High 5 cells at 0.8-1 × 10^6^ cells/mL concentration were infected by adding 1% of virus. Expression was stopped 72 to 96h after the day of proliferation arrest and cells were harvested by centrifugation (1000 g, 20 minutes, 4 degrees). The cells were disrupted by sonication for 4 min (5 sec ON, 20 s OFF, 40% amplitude) on ice in lysis buffer (50 mM HEPES pH 8, 500 mM NaCl, 20 mM imidazole, 0.5 mM TCEP and 10 % glycerol) with cOmplete EDTA-free Protease Inhibitor Cocktail (Roche). After lysate centrifugation at 48,384 g for 45 min at 4°C, the soluble fraction was loaded on a HisTrap HP ion affinity chromatography (Cytiva). Bound proteins were subjected to two sequential washes steps using (i) the lysis buffer supplemented by 1 M NaCl and (ii) the lysis buffer supplemented by 50 mM imidazole. Bound proteins were eluted using initial lysis buffer supplemented by 500 mM imidazole. TiLV heterotrimeric complex fractions were pooled and diluted to reach the heparin-loading buffer concentration (50 mM HEPES pH 8, 250 mM NaCl, 0.5 mM TCEP, 5% glycerol). Proteins were loaded on a HiTrap Heparin HP (Cytiva) column, washed using the heparin-loading buffer and eluted using 50 mM HEPES pH 8, 1 M NaCl, 2 mM TCEP, 5% glycerol. The nucleic acid free TiLV heterotrimeric complex fractions (ratio A_260/280_ = 0.6) were then concentrated using Amicon Ultra (30 kDa cut-off), flash frozen in liquid nitrogen, and stored at -80 C for further use.

### Electron microscopy

#### TiLV polymerase in vRNA pre-initiation state (sample 1)

To trap TiLV polymerase bound to the 5’/3’ vRNA promoters, 1.6 µM of TiLV polymerase were mixed with 4.8 µM of the TiLV vRNA loop in the cryo-EM buffer and incubated for 1h at 4 degrees. Before proceeding to grids freezing, the sample was centrifuged for 5 min, 11000 g and kept at 4 C.

#### TiLV polymerase in cRNA pre-initiation state (sample 2)

To trap TiLV polymerase bound to the 5’/3’ cRNA promoters, 1.6 µM of TiLV polymerase were mixed with 4.8 µM of the TiLV cRNA loop in the cryo-EM buffer and incubated for 1h at 4 degrees. Before proceeding to grids freezing, the sample was centrifuged for 5 min, 11000 g and kept at 4 C.

#### TiLV polymerase in vRNA initiation state (sample 3)

To trap TiLV polymerase in the vRNA initiation state, 1.6 µM of TiLV polymerase were mixed with an equimolar amount of TiLV vRNA loop in a cryo-EM buffer (50 mM HEPES pH 8, 150 mM NaCl, 2 mM TCEP) supplemented with 100 µM CTP and 10 mM MgCl_2_, incubated for 1h at 4 degrees. Before proceeding to grids freezing, the sample was centrifuged for 5 min, 11000 g and kept at 4 C.

#### TiLV polymerase in elongation state (sample 4)

To trap TiLV polymerase in an early-elongation state, 1.6 µM of TiLV polymerase were mixed with 1.3 µM of the TiLV vRNA loop and 16 µM of capped RNA primer 13-mer in the cryo-EM buffer supplemented with 100 µM ATP, 100 µM UTP, 100 µM CpNHpp (Jena

Bioscience) and 10 mM MgCl_2_. The mix was incubated for 4 h at room temperature. Before proceeding to grids freezing, the sample was centrifuged for 5 min, 11000 g and kept at 4 C.

#### Grid preparation and data collection

For grids preparation, 1.5 µl of sample was applied on each sides of glow discharged (PELCO easiGlow™ Glow Discharge Cleaning System: 45 sec, 30 mA, 0.45 mBar) or plasma cleaned (Fischione 1070 Plasma Cleaner: 1 min 10, 90% oxygen, 10% argon) grids (UltrAufoil 1.2/1.3, Au 300). Excess solution was blotted for 3 to 5 s, blot force 0, 100% humidity at 4 degrees with a Vitrobot Mark IV (ThermoFisher) before plunge freezing in liquid ethane.

Automated data collection of the sample 1 was performed on a 200 kV Glacios cryo-TEM microscope (ThermoFisher) equipped with a K2 direct electron detector (Gatan) using SerialEM^37^. Coma and astigmatism correction were performed on a carbon grid. Movies of 40 frames were recorded in counting mode at a x36,000 magnification giving a pixel size of 1.1 Å with defocus ranging from −0.8 to −2.0 μm. Total exposure dose was ∼40 e^−^/Å2.

Automated data collection of the sample 2, 3, 4 were performed on a Titan Krios G3 (Thermo Fisher) operated at 300 kV equipped with a K3 (Gatan) direct electron detector camera and a BioQuantum energy filter, using EPU. Coma and astigmatism correction were performed on a carbon grid. Micrographs were recorded in counting mode at a ×105,000 magnification giving a pixel size of 0.84 Å with defocus ranging from −0.8 to −2.0 µm. For sample 2 and 3, gain-normalized movies of 50 frames were collected with a total exposure of ∼50 e^−^/Å^2^. For sample 4, gain-normalized movies of 40 frames were collected with a total exposure of ∼40 e^−^/Å^2^.

### Image processing

For each collected dataset, movie drift correction was performed using Relion’s Motioncor implementation, with 5x5 patch for K2 movies and 7x5 patch for K3 movies, using all movie frames^38^. For images collected on the Glacios cryo-TEM (ThermoFisher), both gain reference and camera defect corrections were applied. All additional initial image processing steps were performed in CryoSPARC v3.3^39^. CTF parameters were determined using “Patch CTF estimation”, realigned micrographs were then inspected and low-quality images were manually discarded. To obtain an initial 3D reconstruction of TiLV polymerase, particles were picked on few hundreds micrographs using a circular blob with a diameter ranging from 90 to 140 Å. Particles were extracted, 2D classified and subjected to an “ab-initio reconstruction” job. The best initial model was further 3D refined and used to prepare 2D templates. For each dataset, particles were then picked using the template picker and extracted from dose-weighted micrographs using a box size of 220 x 220 pixels^2^ for the Glacios (ThermoFisher) dataset and 300 x 300 pixels^2^ for the Titan Krios (ThermoFisher) datasets.

The same image processing approach was used to separate the different TiLV polymerase conformation. Successive 2D classifications were used to eliminate particles displaying poor structural features. All remaining particles were then transferred to Relion 4.0. For each dataset, particles were divided in subset of 300k to 500k particles and subjected to multiple 3D classification with coarse image-alignment sampling using a circular mask of 140 Å. For each similar TiLV polymerase conformation, particles were grouped and subjected to masked 3D refinement further followed by multiple 3D classification without alignment or using local angular searches. Once particles were properly classified, Bayesian polishing was performed and shiny particles subjected to a last masked 3D refinement. Post-processing was performed in Relion using an automatically or manually determined B-factor. For each final map, reported global resolution is based on the FSC 0.143 cut-off criteria and local resolution variations were also estimated in Relion.

For detailed image processing information regarding each data collection, please refer to Supplementary Information 1-4.

### Model building and refinement

Model building of the full-length TiLV polymerase and associated RNAs was performed *de novo* in COOT^40^. The Robetta^41^ prediction of the structure of the polymerase core of segment 1 was used as a starting point. Models were refined using Phenix real-space refinement^42^ with Ramachandran restraints. Atomic model validation was performed using Molprobity^43^ as implemented in Phenix. Model resolution according to cryo-EM map was estimated at the 0.5 FSC cutoff. Figures were generated using ChimeraX^44^. Electrostatic potential was calculated using PDB2PQR^45^ and APBS^46^.

### *In vitro* polymerase activity assays

#### PA-endo cleavage activity

Synthetic RNAs (Integrated DNA technologies) (i) TiLV 5’ vRNA end 1-15 (5’ - pGCA AAU CUU UCU CAC – 3’) and (ii) TiLV 3’ vRNA end 1-15 (5’ - GUG AGU AAA AUU UGG – 3’) promoters were mixed with 1 µM TiLV polymerase in a 1:1:1 molar ratio. The nuclease activity of TiLV PA-ENDO-like domain was then tested by incubating TiLV polymerase bound or not to the 5’/3’ vRNA promoters with a 41-mer U-rich RNA (5’ – GGC CAU CCU GUU UUU UUU CCC UUU UUU UUU UUC UUU UUU UU) at 2 µM in presence of multiple divalent metal ions (Mn^2+^, Mg^2+^, Zn^2+^, Co^2+^, Ca^2+^) at 1 mM final concentration in a final reaction buffer composed of 50 mM HEPES pH 8, 150 mM NaCl, 2 mM TCEP. Reactions were realized at room temperature for 1h and stopped by adding 2x RNA loading buffer (95% formamide, 1 mM EDTA, 0.025% SDS, 0.025% bromophenol blue, 0.01% xylene cyanol). Samples were heated 5 min at 95 degrees and analysed on a 7 M urea 15% acrylamide gel further stained with SYBR Gold (ThermoFisher). As a positive control, a similar reaction was setup with the A/H7N9 influenza polymerase activated using both 5’ vRNA 1-14 (5’- pAGU AGA AAC AAG GC-3’) and 3’ vRNA 1-6 (5’ –UCU GCU-3’) promoters, in the same ratio as TiLV polymerase reactions, using 1 mM Mn^2+^ in a buffer containing 50 mM HEPES pH 8, 150 mM NaCl, 2 mM TCEP. Reaction was done at 30 degrees for 1h and stopped by adding 2x RNA loading buffer. Sample was heated 5 min at 95 degrees and analysed on a 7M urea 15% acrylamide gel further stained with SYBR Gold (ThermoFisher).

The Decade Markers System (Ambion) was used as a molecular weight ladder.

#### PB2-CBD cap binding assay

TiLV polymerase and A/H7N9 influenza polymerase (positive control) were respectively mixed with TiLV vRNA loop and H7N9 vRNA loop (1:1 molar ratio) in a loading buffer (50 mM HEPES pH 8, 150 mM NaCl, 2 mM TCEP). Polymerases-RNA mixes were then incubated with immobilized γ-Aminophenyl-m7GTP (C10-spacer) beads (Jena Bioscience) for 2h at 4 degrees, washed extensively with the loading buffer and eluted with the loading buffer supplemented with 1 mM m^7^GTP (Jena Bioscience). Input, washed, eluted fractions and what remains bound to the beads were analysed on 4-20% Tris-glycine gel (ThermoFisher) stained with Commassie blue.

#### PB1 RdRp activity

Synthetic RNAs were made by Integrated DNA technologies. Capped RNAs were made by Mathieu Noel and Francoise Debart (Institut des Biomolécules Max Mousseron, Montpellier, France).

TiLV 40-mer vRNA loop, enclosing the 20 first and 20 last nucleotides of segment 9 were used as a model vRNA (5’-pGCA AAU CUU UCU CAC GUC CUG ACU UGU GAG UAA AAU UUG G -3’).

TiLV 40-mer cRNA loop, enclosing the 20 first and 20 last nucleotides of the complement of segment 9, were used as a model cRNA (5’-pCCA AAU UUU ACU CAC AAG UCA GGA CGU GAG AAA GAU UUG C -3’).

Synthetic uncapped 9-mer (5’-UAU AAU ACC -3’) and 10-mer (5’- UAU AAU ACC A - 3’) RNAs and capped 11-mer (5’- m^7^GTP – AAU AUA AUA CC -3’), 12-mer (5’- m7GTP – GAA UAU AAU ACC -3’) and 13-mer (5’- m7GTP – AGA AUA UAA UAC C -3’) RNAs ending by …CC-3’ were used as primers for *in vitro* activity assays.

For nucleotide incorporation activity assays, 0.8 μM TiLV polymerase were mixed with 1 μM TiLV vRNA loop and 8 μM capped primers. Reactions were launched by adding all NTPs or ATP/UTP or ATP/UTP/CpNHpp (100 μM / NTPs) with MgCl_2_, 0.75 μCi/ml α-^32^P ATP in a reaction buffer (50 mM HEPES pH 8, 150 mM NaCl, 2 mM TCEP, 10 mM MgCl_2_) for 4h at 24 C. Reaction temperature and magnesium concentration used were found by optimization (data not shown). Decade Markers System (Ambion) was used as molecular weight ladder. Reactions were stopped by adding 2x RNA loading dye, heating 5 min at 95 C and immediately loaded on a denaturing 20% acrylamide 7 M urea gel. The gel was exposed on a storage phosphor screen and read with an Amersham Typhoon scanner (Cytiva).

#### Data availability

The data that support this study are available from the corresponding authors upon reasonable request. The coordinates and EM maps generated in this study have been deposited in the Protein Data Bank and the Electron Microscopy Data Bank. See Table 1. All uncropped gels are presented in a source data file.

## Acknowledgments

We especially thank Guy Schoehn and Eleftherios Zarkadas for supporting our access to the IBS Glacios. We acknowledge the European Synchrotron Radiation Facility and the Partnership for Structural Biology (PSB) for access to the Titan Krios CM01, and especially Eaazhisai Kandiah for setting up the multiple high-quality data collections. We thank Wojtek Galej, Sarah Schneider and Romain Linares for access to the EMBL Grenoble Glacios and Aymeric Peuch for support using the joint EMBL-IBS computer cluster. We thank Mathieu Noel and Francoise Debart (Institut des Biomolécules Max Mousseron, Montpellier, France) for synthesis of capped RNAs. This work used the platforms at the Grenoble Instruct-ERIC Center (ISBG; UMS 3518 CNRS CEA-UGA-EMBL) with support from the French Infrastructure for Integrated Structural Biology (FRISBI, ANR-10-INSB-05-02) and GRAL, a project of the University Grenoble Alpes graduate school (Ecoles Universitaires de Recherche) CBH-EUR-GS (ANR-17-EURE-0003) within the Grenoble Partnership for Structural Biology. The IBS Electron Microscope facility is supported by the Auvergne Rhône-Alpes Region, the Fonds Feder, the Fondation pour la Recherche Médicale and GIS-IBiSA. The work was partly supported by ANR grant ANR-18-CE11-0028 to SC.

