## Supplemental Tables and Figures for "Structural and functional analysis of the minimal orthomyxovirus-like polymerase of Tilapia Lake Virus from the highly diverged *Amnoonviridae* family"

**Extended Data Legends.**

**Extended data Table 1. Data Collection and model statistics for the twelve cryo-EM structures of TiLV polymerase.**

**Extended data Fig. 1. Molecular biology and biochemistry strategies applied to find other segment proteins interacting with TiLV segment 1 (putative PB1 subunit)**.

**a.** Strategy applied to clone and express the ten TiLV viral proteins from one plasmid. TiLV segment 1 is coloured in blue, segment 2 in orange and segment 3 in green. Symbols are defined at the top right. The psBIG2ab digestion is in agreement with the simulation according to the agarose gel.

**b.** SDS-PAGE analysis of TiLV polymerase following an initial immobilized metal affinity chromatography (IMAC) purification step, with PB1 subunit bearing a C-terminal (Cter) poly-histidine purification tag (his-tag). The molecular ladder (MW) is on the left. “SNP” corresponds to the total fraction. “SN” to the soluble fraction. “FT” to the flow through. “W” to the wash. “E” to the elution. Deduced TiLV segments are annotated based on their molecular weight.

**c.** SDS-PAGE analysis of TiLV polymerase following a second heparin affinity purification step, with PB1 subunit bearing a C-terminal (Cter) poly-histidine purification tag (his-tag). “IN” corresponds to the input. “FT” to the flow through. “E” to the elution. Deduced TiLV segments are annotated based on their molecular weight. Stoichiometry and degradation problems of TiLV segment 2 are visible necessitating the relocation of the his-tag to TiLV segment 2 C-terminus (Cter).

**Extended data Fig. 2. Comparison of TiLV, Thogoto and influenza polymerase PA-N domains.**

**a-c**. PA ENDO or ENDO-like domains of TiLV, Thogoto and Influenza polymerases. Domains are aligned based on their conserved β-sheets, and coloured from the N-terminus (Nter; light green) to the C-terminus (Cter; dark green). Overall and close-up views of each catalytic (or putative) active site are shown, and corresponding residues are displayed. PDB IDs are indicated.

**Extended data Fig. 3. Comparison of TiLV, SFTSV, Thogoto and influenza polymerase PB2 cap-binding domains.**

**a-e**. PB2 CBD or CBD-like domains are shown from TiLV polymerase, SFTSV-L (inhibited), SFTSV L (cap-bound), Thogoto and influenza polymerases. Domains were aligned based on their cap-binding site and are coloured from the N- (Nter; light orange) to C-terminus (Cter; dark orange). TiLV polymerase and SFTSV-L PB2 link domains are coloured in magenta. $m^7$GTP atoms are displayed as spheres (overall view) or sticks (zoom view) and coloured in purple. Close-up views of each cap-binding (or putative) site are shown, and corresponding residues are displayed. Hydrogen bonds are represented as grey dotted lines. PDB IDs are indicated.

**Extended data Fig. 4. TiLV polymerase promoter binding mode A.**

**a.** Schematic summary of all the interactions between TiLV polymerase and the promoter bound in mode A. The 5' and 3' vRNA ends are respectively in pink and gold. Nucleotide differences between vRNA to cRNA are annotated in red. Flexible nucleotides are in transparent. Water molecules are red spheres and magnesium ions ($Mg^{2+}$) green spheres. The 3' end nucleotides are numbered from the 3' to 5'. Interacting residues are coloured according to subunit, green for PA, blue for PB1 and dark red for PB2-N. In mode A, the 3' end is in the RdRp active site. In the vRNA initiation state, CTP (dark grey) is in the position +1 of the active site.

**b.** Extracted RNA density from TiLV polymerase in vRNA pre-initiation state (mode A) cryo-EM map. The +1 active site position is indicated by a dotted line.

**c.** Extracted RNA density from TiLV polymerase in cRNA pre-initiation state (mode A) cryo-EM map. The v to cRNA nucleotide differences are labelled in red. The +1 position is indicated by a dotted line.

**d.** Extracted RNA density from TiLV polymerase in vRNA initiation state cryo-EM map. The +1 position is indicated by a dotted line, which encompasses the incoming CTP.

**Extended data Fig. 5. TiLV polymerase promoter binding mode B.**

**a.** Schematic summary of all the interactions between TiLV polymerase and the promoter bound in mode B. The 5' and 3' vRNA ends are respectively in pink and gold. Nucleotide differences between vRNA to cRNA are annotated in red. Flexible nucleotides are transparent. Water molecules are shown as red spheres. The 3' end nucleotides are numbered from the 3' to 5'. Interacting residues are coloured according to subunit, green for PA, blue for PB1 and dark red for PB2-N. In the closed core conformation, the 3' end nts A7-G15 are not visible, whereas in the open core conformation, they are visible and interact with specific residues that are surrounded by dotted rectangles.

**b.** Extracted RNA density from TiLV polymerase in vRNA pre-initiation state (mode B, close core) cryo-EM map.

**c.** Extracted RNA density from TiLV polymerase in vRNA pre-initiation state (mode B, open core) cryo-EM map.

**d.** Extracted RNA density from TiLV polymerase in cRNA pre-initiation state (mode B, close core) cryo-EM map. The v to cRNA nucleotide differences are coloured in red.

**e.** Comparison of the distal duplex orientation in mode A and B. TiLV polymerase PA, PB1 and PB2 subunits are shown as surfaces and respectively coloured in green, light blue and red. The 5' and 3' vRNA ends are respectively in pink and gold. Flexible nucleotides are represented as dotted lines.

**Extended data Fig. 6. Comparison of 5' hook structures from viral polymerases belonging to the *Articulavirales* and the *Bunyavirales* orders.**

**a-e**. 5' vRNA hooks are shown for TILV, Influenza, Lassa virus (LASV), La Crosse virus (LACV), Hantan virus (HTNV) and severe fever with thrombocytopenia syndrome virus (SFTSV). Nucleotides are numbered and sequentially coloured from the 5' end (light pink) to the 3' end (dark pink). For TiLV flexible nucleotides are indicated by a dotted line. PDB IDs are indicated.

**Extended data Fig. 7. TiLV polymerase, from pre-initiation to elongation state focussing on the active site.**

**a.** Close-up view on TiLV polymerase active site in vRNA pre-initiation state mode A. The 3' vRNA end is coloured in gold and nucleotides are numbered from the 3' to 5', with G1 and G2 being in the respective -1/+1 active site positions. RdRp motifs A, B, C, D, and F are respectively coloured in dark turquoise, purple, grey, light green, and blue. Key residues are displayed. Hydrogen bonds are represented as grey dotted lines. The +1 active site and the NTP entry channel are indicated with arrows. Schematic of the overall RNA conformation is shown on the right with the +1 active site position indicated with a dotted rectangle. Flexible nucleotides are in italic.

**b.** Close-up view on TiLV polymerase active site in cRNA pre-initiation state mode A. The 3' cRNA end is coloured in gold and nucleotides are numbered from the 3' to the 5', with C1 and G2 being in the respective -1/+1 active site position. TiLV RdRp motifs are coloured as in **a**. Water molecules and magnesium ions ($Mg^{2+}$) are respectively represented as red and green spheres. Hydrogen bonds are represented as grey dotted lines. Schematic of the overall RNA conformation is shown on the right with the +1 active site position indicated with a dotted rectangle. Nucleotide differences between the vRNA and the cRNA are coloured in red. The +1 active site position is indicated with a dotted rectangle. Flexible nucleotides are in italic.

**c.** Close-up view on TiLV polymerase active site in vRNA initiation state. TiLV RdRp motifs and RNA are coloured as in **a**. Water molecules and magnesium ions ($Mg^{2+}$) are respectively represented as red and green spheres. Hydrogen bonds are represented as grey dotted lines. The incoming CTP in the +1 active site position is coloured in dark grey. Schematic of the overall RNA conformation is shown on the right with the +1 active site position indicated with a dotted rectangle. Flexible nucleotides are in italic.

**d.** Close-up view on TiLV polymerase active site in vRNA stalled elongation state. TiLV RdRp motifs and RNA are coloured as in **a**. Water molecules and magnesium ions ($Mg^{2+}$) are respectively represented as red and green spheres. Hydrogen bonds are represented as grey dotted lines. The incoming CpNHpp in the +1 active site position is coloured in dark grey. For more clarity, only the last incorporated nucleotide from the product is shown, and coloured in cyan. Schematic of the overall RNA conformation is shown on the right with the +1 active site position and the CpNHpp indicated with a dotted rectangle. The capped primer ending by ...CC-3' (13-mer) is coloured in magenta and the incorporated nucleotides are coloured in cyan. Flexible nucleotides are in italic.

**Extended data Fig. 8. TiLV polymerase in elongation states.**

**a.** Extracted RNA density from TiLV polymerase in vRNA elongation state cryo-EM map. The 5' vRNA end is coloured in pink. The 3' vRNA end is coloured in gold. The capped primer

(13-mer) is coloured in magenta and the incorporated nucleotides are coloured in blue. The CpNHpp in the +1 active site position is coloured in dark grey. Flexible nucleotides are represented as dotted lines.

**b.** Extracted RNA density from TiLV polymerase in vRNA elongation state cryo-EM map (distal duplex). The 3' vRNA end excess is able to bind the secondary site, reforming a distal duplex with the 5' vRNA end. The RNA colour code is identical to **a**.

**c.** Schematic of the overall RNA conformation. The +1 active site position and the CpNHpp are indicated with a dotted rectangle. The capped primer ending by ...CC-3' (13-mer) is coloured in magenta and the incorporated nucleotides are coloured in cyan. Flexible nucleotides are in italic.

Schematic summary of all interactions between TiLV polymerase, the transcription-like product and the 3' vRNA end template. The RNA colour code is identical to **a**. The 3' vRNA end nucleotides are numbered from the 3' to the 5'. Flexible nucleotides are represented as dotted line. Water molecules and magnesium ions ($Mg^{2+}$) are respectively represented as red and green spheres. Interacting residues are coloured according to their TiLV polymerase subunits, green for PA, blue for PB1 and dark red for PB2-N.

**Extended data Fig. 9. Comparison of TiLV polymerase initiation and elongation states with FluPol and LACV-L.**

**a.** Structural comparison of TiLV polymerase in initiation state with FluPol in pre-initiation state and LACV-L in initiation state. PA is coloured in green, PB1 in light blue and PB2-N in dark red. 5' vRNA ends are coloured in pink. 3' vRNA ends are coloured in gold. FluPol priming loop and LACV-L template exit plug are coloured in red and residues are shown as spheres. The template entry, template exit, and the secondary site are indicated. PDB IDs are indicated.

**b.** Structural comparison of TiLV polymerase, FluPol and LACV-L in their respective early-elongation state. Domains and RNAs are shown and coloured as in **a**. Upon elongation, FluPol priming loop and LACV-L template exit plug are extruded. The template entry, template exit, and the secondary site are indicated. PDB IDs are indicated.

**c.** Structural comparison of TiLV polymerase in early-elongation with distal duplex reformation, FluPol in pre-termination state and LACV-L in late-elongation state. Domains and RNAs are shown and coloured as in **a**. Upon late-elongation, the 3' end is able to bind back to the secondary site. Flexible nucleotides are shown as dotted line. The template entry, template exit, and the secondary site are indicated.PDB IDs are indicated.

**Extended data Fig. 10. Alternative configurations of TiLV polymerase.**

**a.** TiLV polymerase transcriptase conformation. Close-up view of PA ENDO-like, PB2-N, and PB2-C domains coloured as in **Fig. 1d**. TiLV polymerase ENDO-like and CBD-like residues analogous to those in the respective FluPol ENDO active site and FluPol cap-binding site are shown as spheres. The PB1 subunit is displayed as a blue surface.

**b.** TiLV polymerase replicase conformation, annotated as in **a**. TiLV PB2 627 domain is flexible in this conformation.

**c.** The transcriptase to replicase transition. Domains are coloured as in **a**. To switch from transcriptase to replicase, PA ENDO-like domain rotates ~160 degrees and PB2 NLS domain translocates to interact with it. PB2 CBD-like domain rotates ~180 degrees and packs against TiLV polymerase PB1 subunit (left). In the transcriptase conformation, TiLV polymerase PA ENDO-like and PB2 CBD-like residues, analogous to the respective FluPol PA ENDO active site and FluPol PB2 cap-binding site, are 32 Å apart. In the replicase conformation the corresponding residues are 50 Å apart (right).

**d.** Close-up view of the TiLV polymerase PA ENDO-like / PB2 NLS domain interaction. Domains are coloured as in **a**. Hydrogen bonds are shown as grey dotted lines. Key interacting residues are shown as sticks.

**e.** In full-length structures, compared to core-only structures, packing of TiLV polymerase 627-like domain against the PB1 subunit induces a conformational change that propagates towards the RdRp active site. The PB2 627-like domain is coloured in pink. PB1 subunit is coloured in light blue with the palm domain coloured in orange. The RdRp motif C (285-295) is indicated. The main movements between TiLV polymerase in vRNA initiation state (core only and complete structure) are indicated with arrows. TiLV polymerase structures are aligned with each other using PB1 as reference.

**Extended data Table 1a. Cryo-EM data collection, refinement and validation statistics of TiLV polymerase structures.**

| | vRNA initiation | | cRNA pre-initiation mode A | | cRNA pre-initiation mode B | vRNA pre-initiation mode A |
|---|---|---|---|---|---|---|
| **Structure No.** | 1 | 2 | 3 | 4 | 5 | 6 |
| **Short name** | Full vRNA initiation with CTP | Core vRNA initiation with CTP | Core cRNA pre-initiation mode A | Core with endo cRNA pre-initiation mode A | Closed core with endo cRNA mode B | Core vRNA pre-initiation mode A |
| | PDB ID 8PSN, EMD-17857 | PDB ID 8PSO, EMD-17858 | PDB ID 8PSQ, EMD-17860 | PDB ID 8PT7, EMD-17869 | PDB ID 8PSS, EMD-17861 | PDB ID 8PSU, EMD-17862 |
| **Data collection and processing** | ESRF CM01 ThermoFisher Krios TEM \| Gatan K3 direct electron detector mounted on a Gatan Bioquantum LS/967 energy filter | | ESRF CM01 ThermoFisher Krios TEM \| Gatan K3 direct electron detector mounted on a Gatan Bioquantum LS/967 energy filter | | | ThermoFisher Glacios TEM \| Gatan K2 Summit |
| **Magnification** | 105000 | | 105000 | | | 36000 |
| **Voltage (kV)** | 300 | | 300 | | | 200 |
| **Electron exposure (e–/Å$^2$)** | 51 | | 51 | | | 40 |
| **Defocus range (μm)** | -0.8 / -2.0 | | -0.8 / -2.0 | | | -0.8 / -2.0 |
| **Pixel size (Å)** | 0.84 | | 0.84 | | | 1.1 |
| **Symmetry imposed** | C1 | | C1 | | | C1 |
| **Initial/Final micrographs** | 6000 / 4133 | | 6000 / 3463 | | | 968 / 953 |
| **Final particle images (no.)** | 46595 | 271170 | 107798 | 153101 | 109188 | 67042 |
| **Map resolution (Å)<br>FSC threshold 0.143** | 2.73 | 2.40 | 2.65 | 2.80 | 2.83 | 3.18 |
| **Map resolution range (Å)** | 2.6-3.8 | 2.3-3.1 | 2.6-3.8 | 2.6-3.8 | 2.6-3.8 | 3.2-4.0 |
| **Refinement** | | | | | | |
| **Model resolution (Å)<br>FSC threshold 0.5** | 2.7 | 2.4 | 2.6 | 2.8 | 2.8 | 3.2 |
| **Map sharpening *B* factor (Å$^2$)** | -40 | -54 | -58 | -40 | -40 | -81 |
| **CC (mask)** | 0.8726 | 0.8841 | 0.8868 | 0.8949 | 0.8870 | 0.8198 |
| **Composition<br>Non-H atoms<br>Protein<br>Nucleotides<br>Water<br>Ligands** | <br>11301<br>1361<br>29<br>2<br>3 x Zn 2 x Mg | <br>8264<br>971<br>29<br>44<br>1 x Zn 2 x Mg | <br>8181<br>969<br>28<br>1<br>1 x Zn 1 x Mg | <br>9032<br>1071<br>29<br>0<br>1 x Zn 1 x Mg | <br>8697<br>1068<br>15<br>0<br>1 x Zn 1 x Mg | <br>8212<br>971<br>29<br>0<br>1 x Zn |
| ***B* factors (Å$^2$)<br>Protein<br>Nucleotides<br>Water<br>Ligands** | <br>38.03<br>63.64<br>19.39<br>66.15 | <br>32.03<br>54.21<br>24.31<br>38.48 | <br>36.52<br>65.87<br>72.78<br>23.03 | <br>66.89<br>102.87<br>-<br>62.76 | <br>72.72<br>85.07<br>-<br>65.56 | <br>49.64<br>91.53<br>-<br>72.80 |
| **RMS deviations<br>Bond lengths (Å)<br>Bond angles (°)** | <br>0.002<br>0.519 | <br>0.002<br>0.482 | <br>0.003<br>0.471 | <br>0.002<br>0.436 | <br>0.002<br>0.440 | <br>0.003<br>0.446 |
| **Validation<br>MolProbity score<br>Clashscore<br>Poor rotamers (%)** | <br>1.50<br>3.50<br>2.57 | <br>1.18<br>2.41<br>1.55 | <br>1.14<br>2.79<br>1.19 | <br>1.35<br>3.66<br>1.29 | <br>1.55<br>2.96<br>1.73 | <br>1.22<br>3.46<br>0.0 |
| **Ramachandran<br>Favored (%)<br>Allowed (%)<br>Disallowed (%)** | <br>97.79<br>2.14<br>0.07 | <br>97.93<br>2.07<br>0.0 | <br>97.92<br>1.97<br>0.10 | <br>97.46<br>2.44<br>0.09 | <br>95.85<br>4.15<br>0.0 | <br>97.62<br>2.18<br>0.21 |

#### Extended data Table 1b. Cryo-EM data collection, refinement and validation statistics of TiLV polymerase structures.

| | vRNA elongation | | vRNA mode B | | | Replicase |
|---|---|---|---|---|---|---|
| **Structure No.** | 7 | 8 | 9 | 10 | 11 | 12 |
| **Short name** | Full vRNA elongation with CpNHpp | Full vRNA elongation with CpNHpp and additional mode B promoter | Full with open core vRNA mode B | Open core with rotated endo-NLS vRNA mode B | Closed core with rotated endo-NLS vRNA mode B | Replicase initiation with CpNHpp |
| | PDB ID 8PSX, EMD-17864 | PDB ID 8PSZ, EMD-17865 | PDB ID 8PT2, EMD-17866 | PDB ID 8PTH, EMD-17871 | PDB ID 8PTJ, EMD-17872 | PDB ID 8PT6, EMD-17868 |
| **Data collection and processing** | ESRF CM01 ThermoFisher Krios TEM \| Gatan K3 direct electron detector mounted on a Gatan Bioquantum LS/967 energy filter | | | | | |
| **Magnification** | 105000 | | | | | |
| **Voltage (kV)** | 300 | | | | | |
| **Electron exposure (e–/Å$^2$)** | 51 | | | | | |
| **Defocus range (µm)** | -0.8 / -2.0 | | | | | |
| **Pixel size (Å)** | 0.84 | | | | | |
| **Symmetry imposed** | C1 | | | | | |
| **Final particle images (no.)** | 29892 | 103012 | 80382 | 36587 | 53296 | 80175 |
| **Initial/Final micrographs (no.)** | 6000 / 5831 | | | | | |
| **Map resolution (Å) FSC threshold 0.143** | 2.96 | 2.42 | 2.59 | 2.73 | 2.86 | 2.9 |
| **Map resolution range (Å)** | 2.6-4.2 | 2.3-3.5 | 2.4-4.0 | 2.5-4.1 | 2.5-4.1 | 2.8-4.8 |
| **Refinement** | | | | | | |
| **Model resolution (Å) FSC threshold 0.5** | 2.9 | 2.4 | 2.6 | 2.7 | 2.9 | 3.0 |
| **Map sharpening *B* factor (Å$^2$)** | -40 | -30 | -20 | -20 | -20 | -60 |
| **CC (mask)** | 0.8368 | 0.8803 | 0.8835 | 0.8854 | 0.8938 | 0.8378 |
| **Composition** | | | | | | |
| **Non-H atoms** | 11486 | 11903 | 11190 | 9245 | 9034 | 10397 |
| **Protein residues** | 1357 | 1356 | 1355 | 1104 | 1102 | 1243 |
| **Nucleotides** | 38 | 53 | 27 | 27 | 15 | 29 |
| **Water** | 0 | 109 | 1 | 1 | 2 | 0 |
| **Ligands** | 3 x Zn 2 x Mg | 3 x Zn 2 x Mg | 3 x Zn | 1 x Zn | 1 x Zn | 3 x Zn 2 x Mg |
| ***B* factors (Å$^2$)** | | | | | | |
| **Protein** | 37.62 | 29.60 | 54.98 | 59.26 | 65.11 | 80.08 |
| **Nucleotides** | 39.86 | 44.81 | 98.50 | 82.32 | 72.32 | 95.52 |
| **Water** | - | 16.22 | 28.66 | 36.24 | 44.34 | - |
| **Ligand** | 81.58 | 60.93 | 115.45 | 91.94 | 101.42 | 122.29 |
| **R.m.s. deviations** | | | | | | |
| **Bond lengths (Å)** | 0.003 | 0.009 | 0.002 | 0.002 | 0.004 | 0.003 |
| **Bond angles (°)** | 0.505 | 0.866 | 0.446 | 0.438 | 0.485 | 0.535 |
| **Validation** | | | | | | |
| **MolProbity score** | 1.61 | 1.57 | 1.36 | 1.64 | 1.80 | 1.64 |
| **Clashscore** | 3.37 | 2.52 | 2.40 | 3.02 | 3.52 | 5.67 |
| **Poor rotamers (%)** | 1.71 | 2.06 | 1.37 | 1.88 | 2.82 | 1.59 |
| **Ramachandran** | | | | | | |
| **Favored (%)** | 95.69 | 95.69 | 96.43 | 95.16 | 95.50 | 96.92 |
| **Allowed (%)** | 4.31 | 4.31 | 3.57 | 4.84 | 4.50 | 2.92 |
| **Disallowed (%)** | 0.00 | 0.00 | 0.0 | 0.0 | 0.0 | 0.16 |

### EXTENDED DATA FIGURE 1

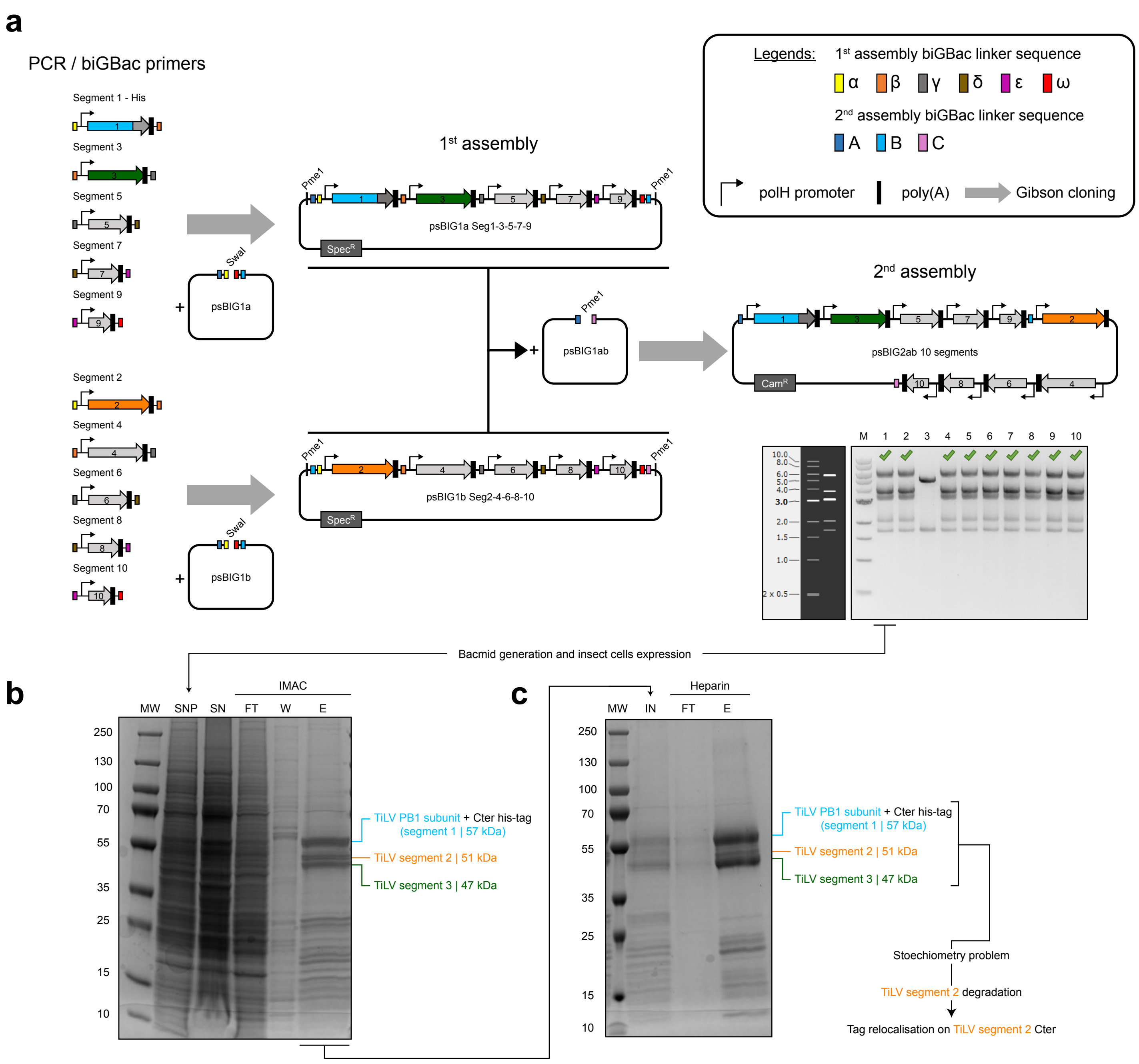

### EXTENDED DATA FIGURE 2

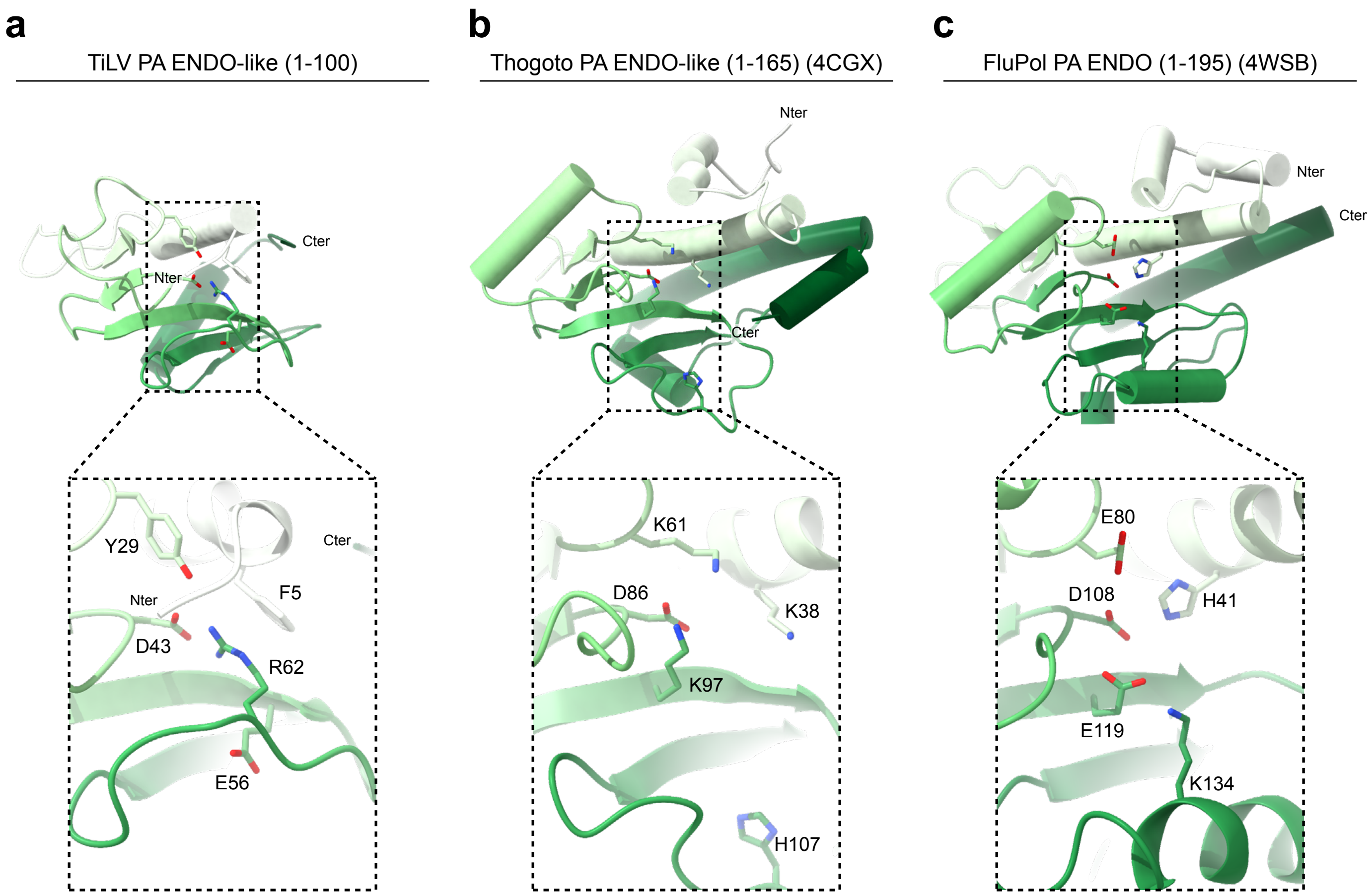

### EXTENDED DATA FIGURE 3

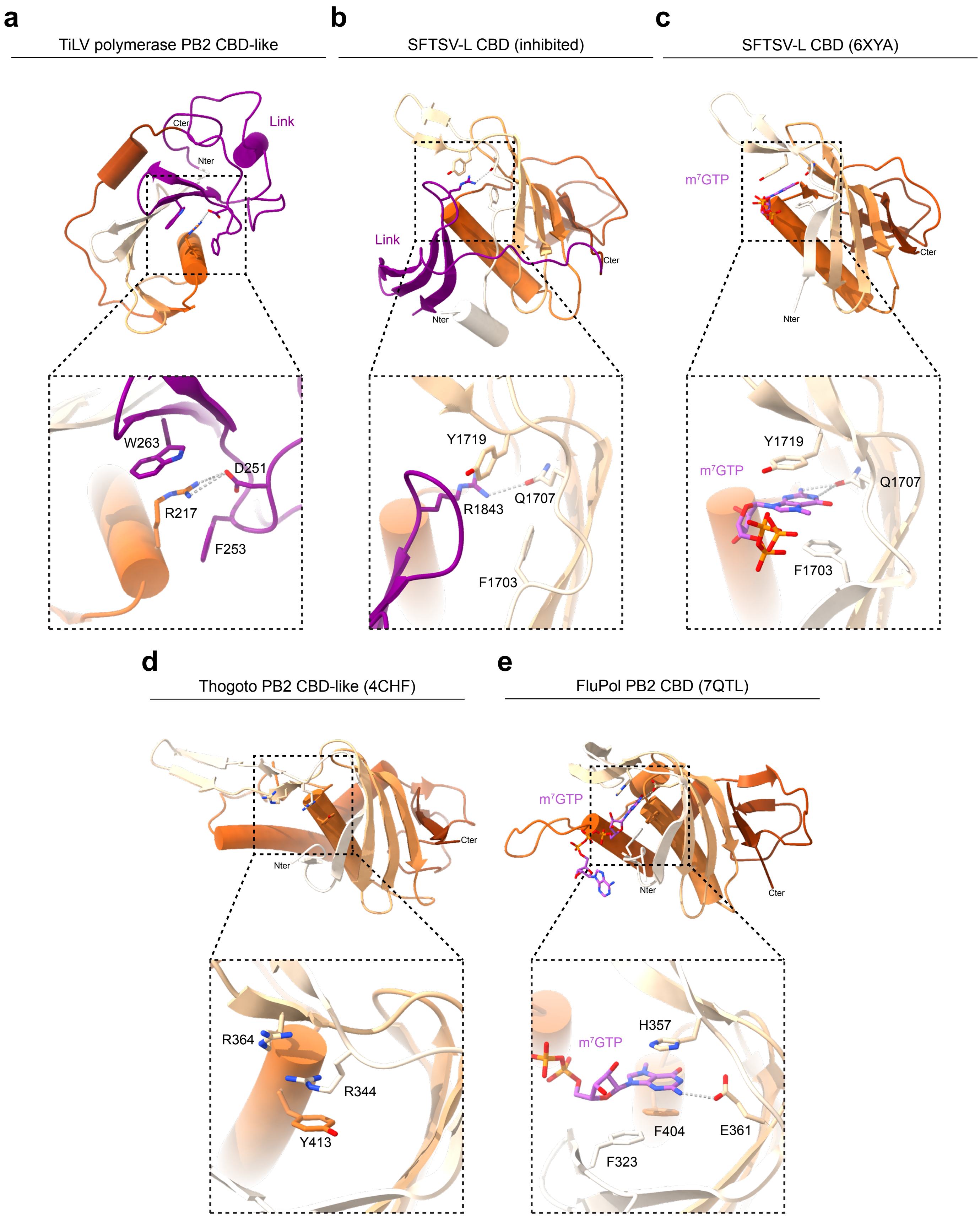

### EXTENDED DATA FIGURE 4

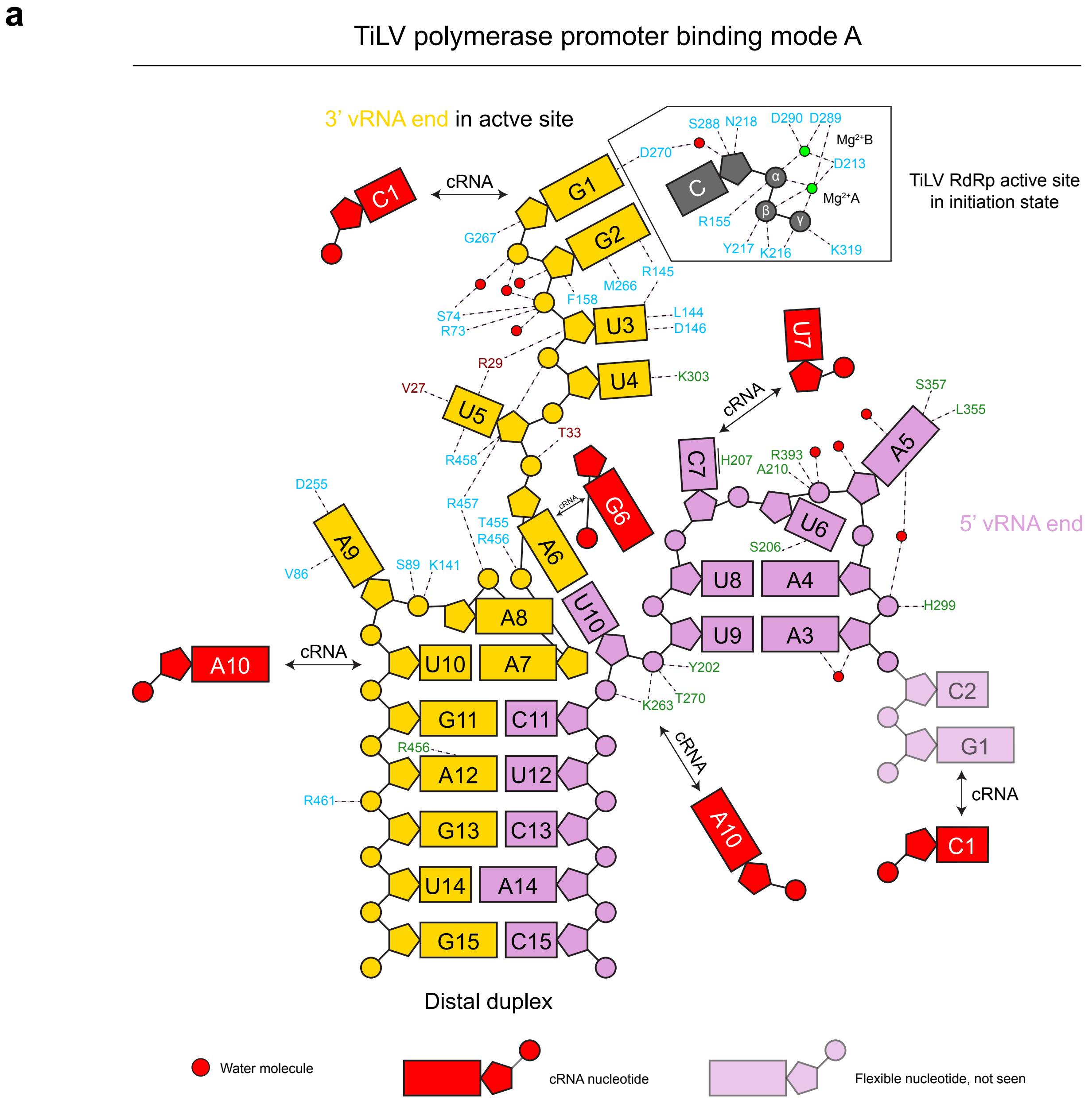


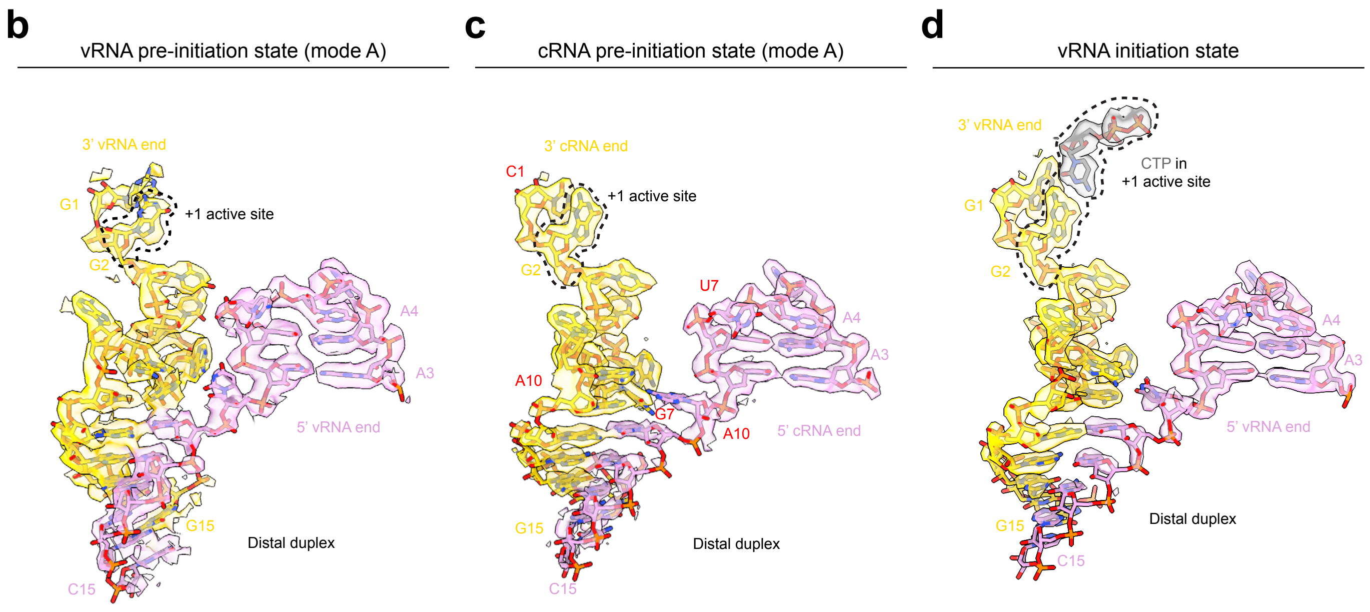

### EXTENDED DATA FIGURE 5

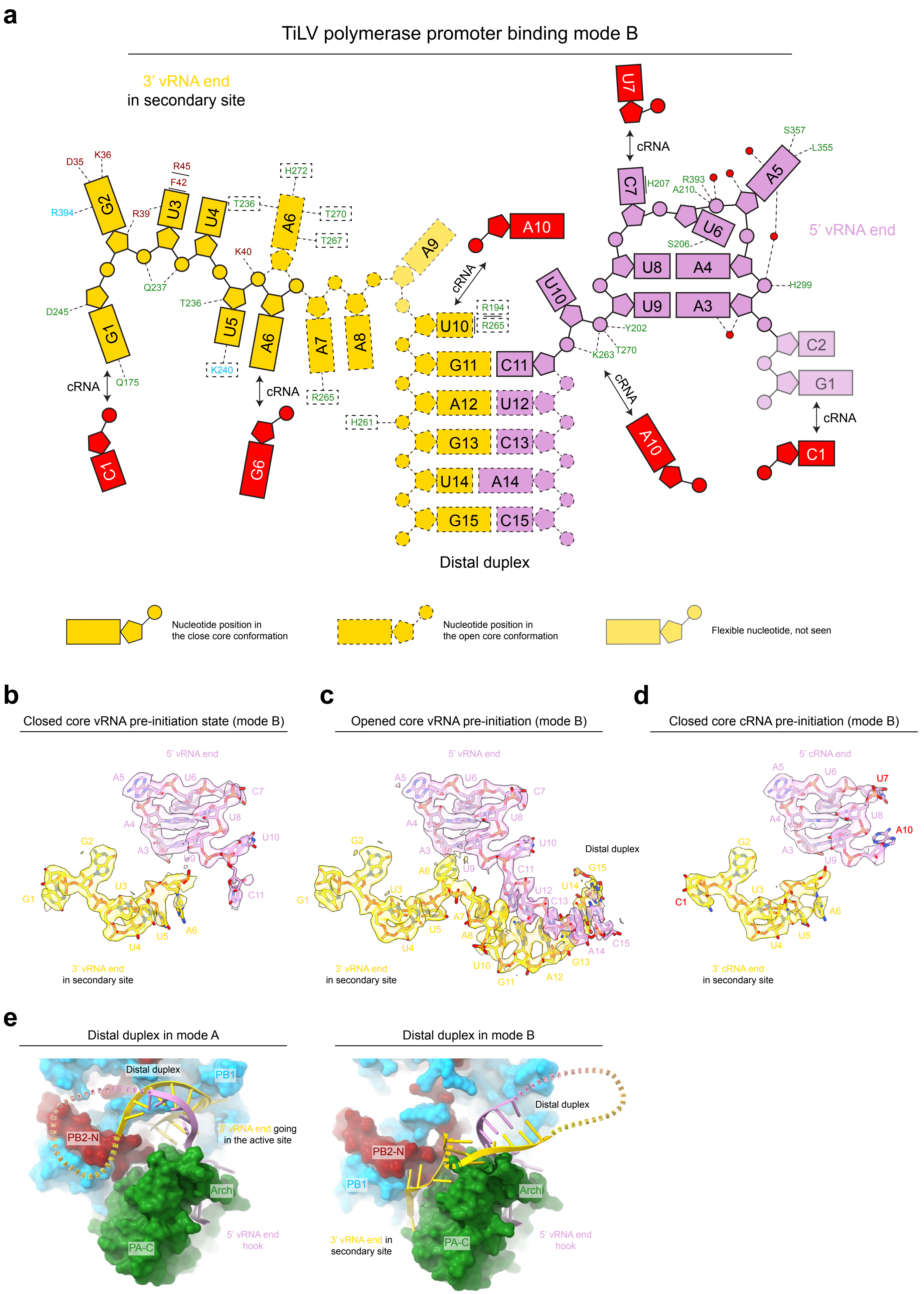

### EXTENDED DATA FIGURE 6

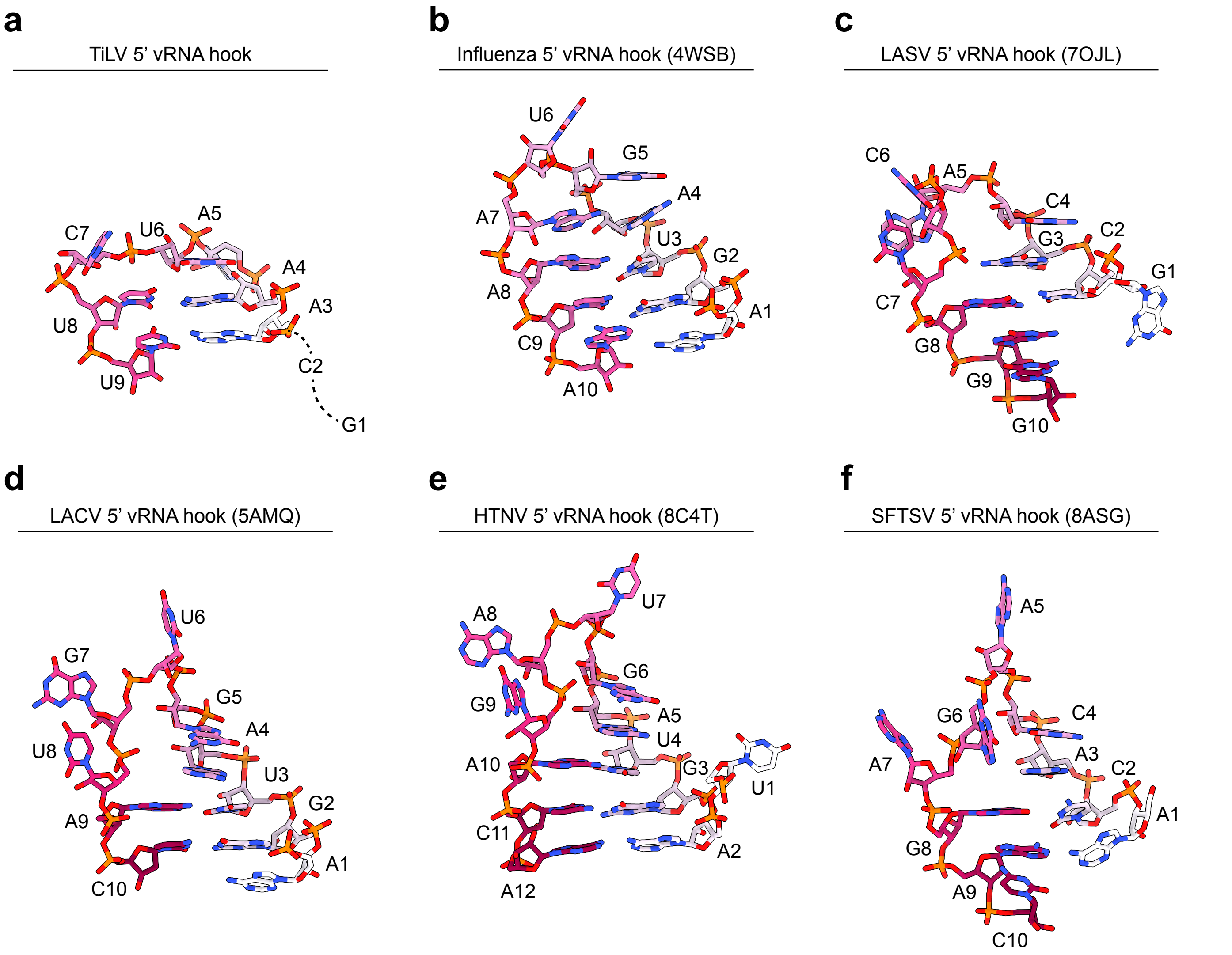

### EXTENDED DATA FIGURE 7

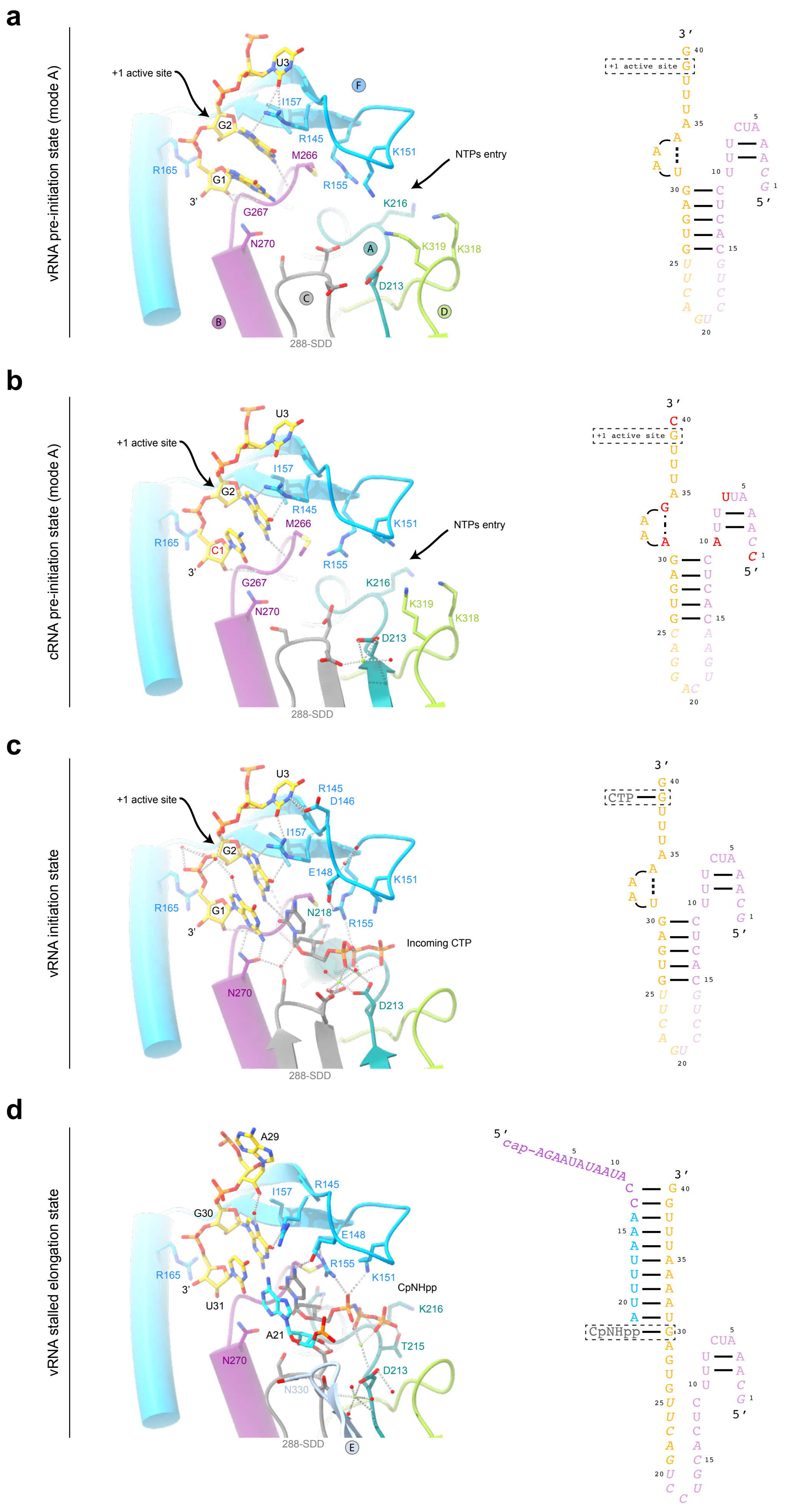

### EXTENDED DATA FIGURE 8

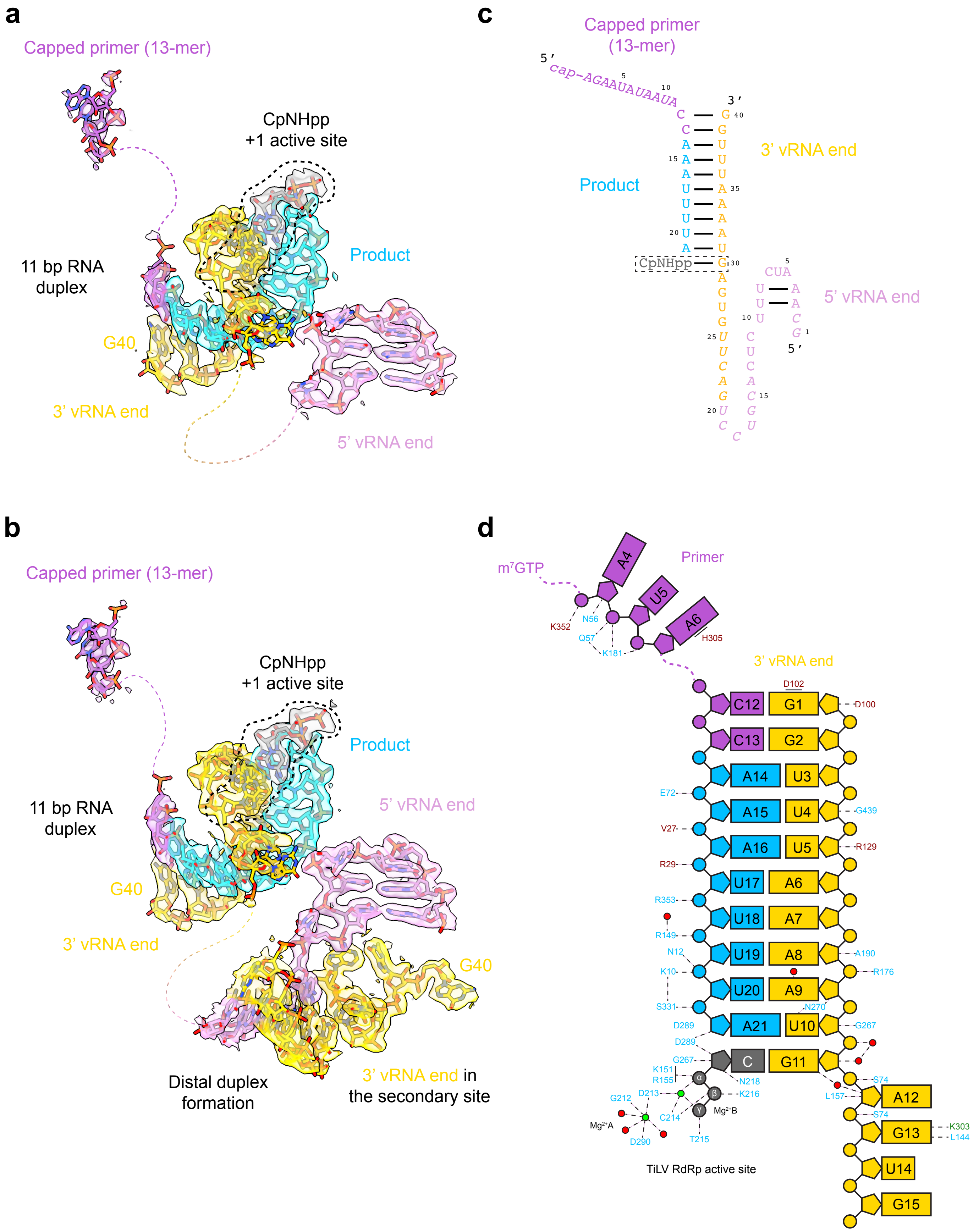

### EXTENDED DATA FIGURE 9

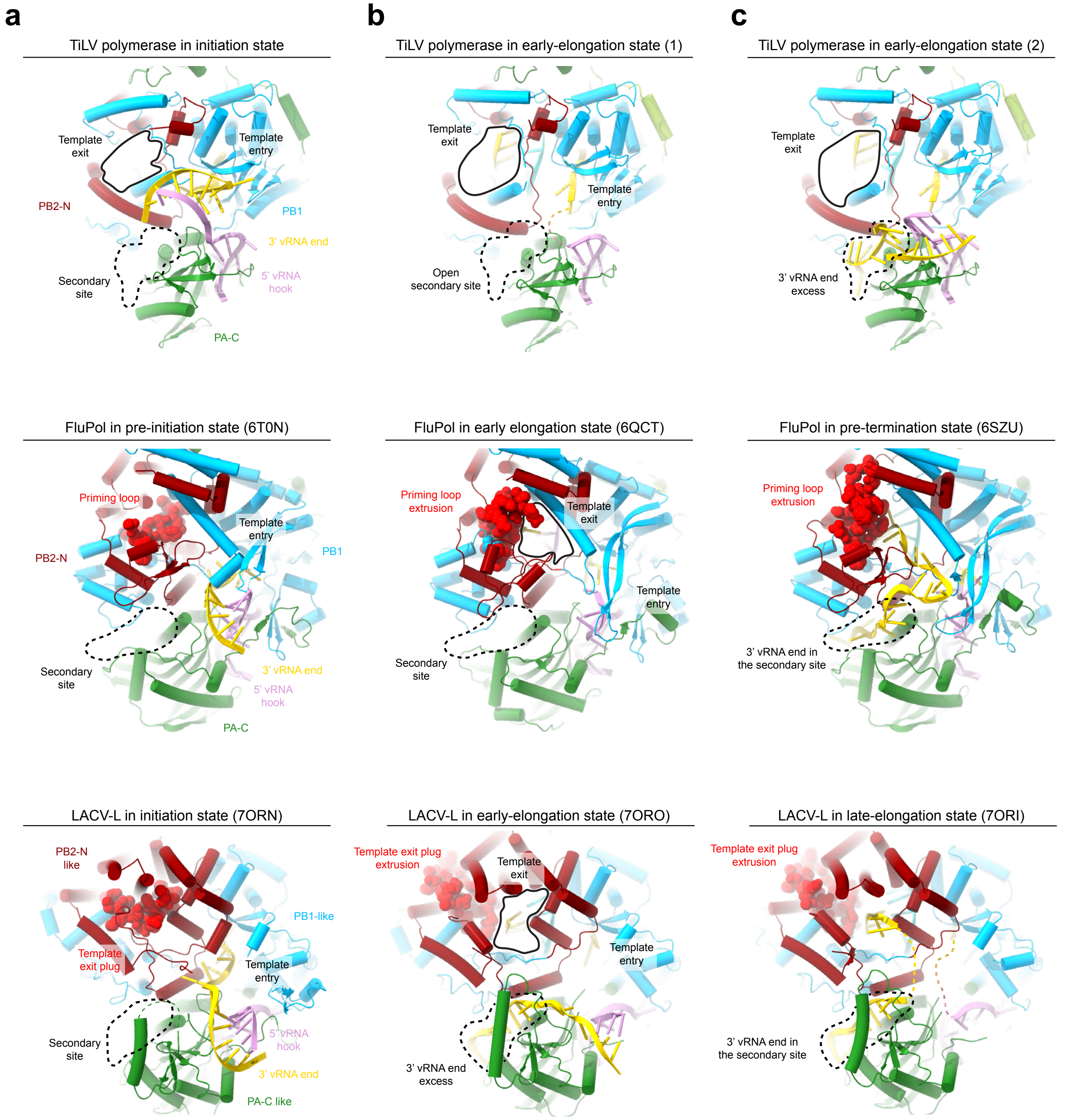

### EXTENDED DATA FIGURE 10

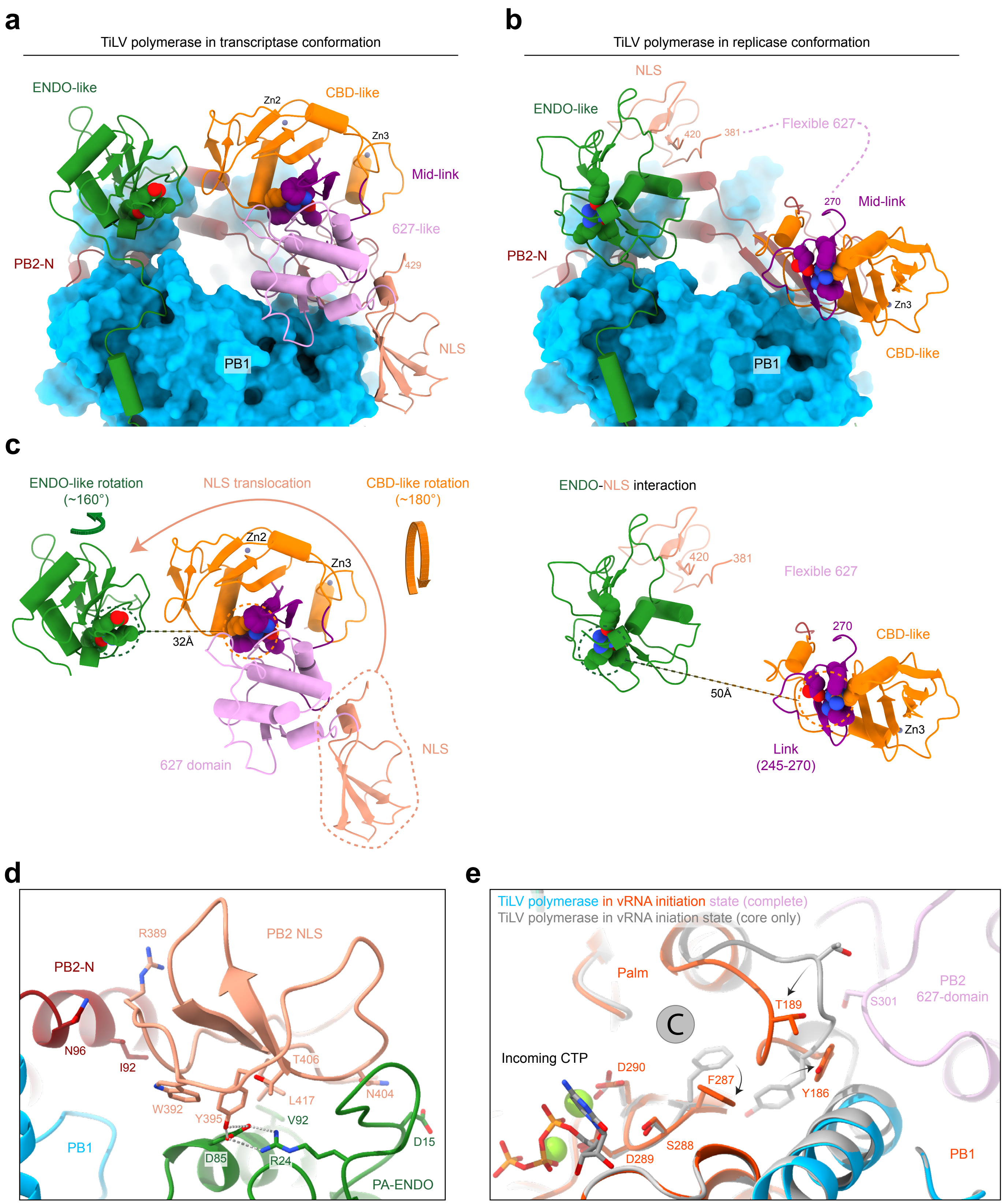
